# Cannabinoid type 2 receptor inhibition enhances the antidepressant and proneurogenic effects of physical exercise after chronic stress

**DOI:** 10.1101/2023.04.24.538087

**Authors:** RS Rodrigues, JB Moreira, SH Vaz, A Barateiro, SL Paulo, JM Mateus, DM Lourenço, FF Ribeiro, E Loureiro-Campos, P Bielefeld, A Fernandes, AM Sebastião, L Pinto, CP Fitzsimons, S Xapelli

## Abstract

Chronic stress is a major risk factor of neuropsychiatric conditions such as depression. Adult hippocampal neurogenesis (AHN) has emerged as a promising target to counteract stress-related disorders given the ability of newborn neurons to facilitate endogenous plasticity. Recent data sheds light on the interaction between cannabinoids and neurotrophic factors underlying the regulation of AHN, with important effects upon cognitive plasticity and emotional flexibility. Since physical exercise (PE) is known to enhance neurotrophin levels, we hypothesized that PE could engage with cannabinoids to influence AHN and that this would result in beneficial effects under stressful conditions. We therefore investigated the actions of modulating cannabinoid type 2 receptors (CB2R), which are devoid of psychotropic effects, in combination with PE in chronically stressed animals. We found that CB2R inhibition, but not CB2R activation, in combination with PE significantly ameliorated stress-evoked emotional changes and cognitive deficits. Importantly, this combined strategy critically shaped stress-induced changes in AHN dynamics, leading to a significant increase in the rates of cell proliferation and differentiation of newborn neurons, and an overall reduction in neuroinflammation. Together, these results show that CB2Rs are crucial regulators of the beneficial effects of PE in countering the effects of chronic stress. Our work emphasizes the importance of understanding the mechanisms behind the actions of cannabinoids and PE and provides a framework for future therapeutic strategies to treat stress-related disorders that capitalize on lifestyle interventions complemented with endocannabinoid pharmacomodulation.

## 1. Introduction

Chronic stress is a major trigger for the development of brain pathologies, particularly major depressive disorder (MDD). MDD is a chronic psychiatric condition that affects more than 300 million people worldwide, a leading cause of disability, and a major contributor to disease burden [1]. Current treatments are still generally inefficient, with frequent relapsing episodes, resulting in a staggering one third of unsuccessfully treated individuals [2]. Therefore, there is an urgent need for new therapeutic avenues focused on effective, long-lasting approaches. Adult neurogenesis, a process whereby new neurons are generated from adult neural stem cells (NSCs) in restricted areas of the adult brain, has emerged as a promising strategy to counter stress-related neuropsychiatric disorders [3]. Specifically, adult hippocampal neurogenesis (AHN), which occurs in the subgranular zone (SGZ) of the hippocampal dentate gyrus (DG), participates in learning and memory as well as emotional and motivational regulation [4]. This process is deleteriously affected by stress [5] and perturbations in AHN dynamics have been associated with MDD pathophysiology [3, 6]. In line with this, human data shows that prolonged exposure to stress results in the reduction of hippocampal volume, dendritic arborization, and AHN [7] whereas resilience is associated with larger DG volumes [8]. Similarly, preclinical studies have found that AHN suppression is involved in processes of susceptibility and maladaptive responses to chronic stress and depressive-like behaviour in rodents, while AHN-enhancing strategies ameliorate these phenotypes [9–11]. Furthermore, the effectiveness of antidepressant therapy was shown to be dependent on AHN [12] while antidepressants were demonstrated to enhance AHN [13], suggesting an interplay between AHN and antidepressant therapy. Additionally, chronic stress and antidepressant therapy were shown to affect, in opposing directions, the expression of endocannabinoids and neurotrophic factors [14, 15], which are known important regulators of AHN [16, 17].

The endocannabinoid system (ECS) has been identified as an important regulator of AHN and stress management [16, 18]. As such, the ECS has been proposed as a key neuromodulatory factor in MDD pathophysiology [19]. Cannabinoids were shown to exert anxiolytic and antidepressant-like effects through hippocampal NSC regulation [20, 21]. Specifically, cannabinoid type 2 receptors (CB2R), which are devoid of psychotropic effects [16], are emerging as crucial players in stress response given their role in the regulation of hippocampal-related plasticity [16, 22, 23]. CB2R activation was shown to exert antidepressant effects [24] while animals lacking CB2Rs displayed exacerbated stress-induced neuroinflammatory responses [25]. Moreover, overexpression of CB2R in mice produced a depression-resistant phenotype with reduced vulnerability to anxiety [26, 27]. Contrastingly, however, chronic inhibition of CB2Rs was found to induce anxiolytic-like effects [28], highlighting the need for further studies addressing the role of CB2Rs in depressive-like behaviours. In human studies, a single CB2R polymorphism was shown to be related with MDD susceptibility, further supporting CB2R therapeutic potential for the treatment of stress-related pathologies [29]. Of note, MDD patients have lower levels of circulating endocannabinoids, suggesting that ECS hypoactivity may play a role in the pathophysiology of the disease [30].

Physical exercise (PE) is known to improve cognitive functions, exert anxiolytic-like effects and regulate hippocampal function and related plasticity mechanisms, namely AHN [31]. Some of the PE-induced beneficial effects on the brain, particularly in the context of MDD, have been attributed to the actions of neurotrophic factors [32, 33]. Indeed, these molecules are known to regulate PE-mediated effects, being important players in plasticity- and pathology-related phenomena like AHN and stress response, respectively [14]. AHN regulation by PE, in which brain-derived neurotrophic factor (BDNF) assumes a preponderant role, has been extensively studied [14, 34]. BDNF critically regulates the survival, dendritic growth, and maturation of newborn SGZ neurons [35, 36]. Remarkably, PE has also been found to inhibit neuroinflammation, a major hallmark in MDD [37].

Recent evidence supports an interaction between PE and the ECS underlying the regulation of AHN [38], with cannabinoids and neurotrophic factors acting as key modulators of cognitive plasticity and mood flexibility [39–41]. In fact, a correlation has been found between the ECS and the mood-enhancing effects of prescribed acute PE in MDD [42]. Additionally, recent evidence from human studies has revealed that a single session of moderate PE influences the levels of endocannabinoids and BDNF, with clear benefits for memory consolidation [43], suggesting a synergistic action of endocannabinoids and BNDF in regulating plasticity-related events. In preclinical studies, this interaction has also been shown, with CB2Rs taking on a preeminent role in regulating BDNF-mediated postnatal neurogenesis [44]. Moreover, CB2R inhibition was found to induce anxiolytic-like effects and dampen BDNF signalling in chronically stressed animals, suggesting a complex interaction between CB2Rs and BDNF in emotional response [45]. However, how CB2Rs may guide the actions of PE in countering the effects of chronic stress has not been addressed. Considering the relationship between PE, neurotrophins and ECS as well as the association of neurotrophins, stress and neurogenesis, we sought to explore whether modulating CB2R activity in combination with PE could revert the neurobiological and behavioural consequences of chronic stress, possibly functioning as a novel antidepressant strategy. We show that reducing CB2R constitutive activity in combination with PE restores chronic stress-induced emotional and cognitive behavioural alterations as well as AHN deficits. Moreover, we found that CB2R inhibition in combination with PE ameliorates chronic stress-induced changes in microglial, astroglial and myelin expression. Hence, these results reveal the potential of combined behavioural and pharmacological strategies for the treatment of depressive disorders and underscore the role of such multimodal approaches for the regulation of AHN and its impact in stress-related pathologies.

## 2. Material and Methods

### 2.1. Ethics and Experimental outline

#### 2.1.1. Animals and Ethical approval

All experiments were performed with male C57Bl6/J mice, aged 14 weeks old, weighing 26-30g, obtained from Charles River Laboratories (Barcelona, Spain). All animals were housed under standard conditions (19–22°C, humidity 55%, 12:12 h light: dark cycle with lights on at 08:00, food and water *ad libitum*), unless stated otherwise. All procedures were conducted in accordance with the European Community (86/609/EEC; 2010/63/EU; 2012/707/EU) and Portuguese (DL 113/2013) legislation for the protection of animals used for scientific purposes. The protocol was approved by the institutional animal welfare body, ORBEA-iMM, and the National competent authority, DGAV (*Direcção Geral de Alimentação e Veterinária*) under the project reference 0421/000/000/2018.

#### 2.1.2. Experimental design and interventional procedures

Animals were randomly assigned to experimental groups after a 1-week habituation and a 1-week handling periods. A dose-response for CB2R ligands (0.5 and 5 mg/kg) was performed considering previous pharmacological ranges [25, 28, 29, 45]. The CB2R agonist HU308 (Tocris, Bristol, UK; abbreviated nomenclature: HU) or the inverse agonist/antagonist AM630 (Tocris, Bristol, UK; abbreviated nomenclature: AM) were administered for 2 weeks, as follows. Unpredictable chronic mild stress (uCMS) was induced as previously described [46–48]. Briefly animals were subjected to an 8-week unpredictable chronic stress paradigm followed by a 2-week milder protocol period during which CB2R ligands were administered in combination with a physical exercise (PE) protocol as previously described [49]. Four weeks before sacrifice, all animals received intraperitoneal (i.p.) injections of BrdU (50 mg/kg; Sigma Aldrich, MO, USA) during 5 consecutive days (twice per day) to evaluate the survival of newborn neurons. Further details are described in supplementary information.

### 2.2. Behavioural analyses

Behavioural tests were performed to assess anxiety- and depressive-like behaviours and cognitive performance after habituation to the experimental room 1h prior to testing in the following order: open field (OF), elevated-plus maze (EPM), novel object recognition (NOR), forced-swimming test (FST) and sucrose splash test (SST). Behavioural testing was carried out using standardised test procedures [50–54], as detailed in supplementary information.

### 2.3. Animal sacrifice, immunostainings and stereology

After an overdose with sodium pentobarbital animals were transcardially perfused with phosphate-buffered saline (PBS). Brains were then extracted and fixed overnight in 4% paraformaldehyde. To evaluate cell proliferation, neuronal differentiation and survival of newborn neurons, particularly discriminating between dorsal (dHip) and ventral (vHip) regions, as well as glia density (i.e., astrocytes, microglia and myelin), brain coronal cryosections (40 µm thick) collected onto a suspension solution were incubated at 4°C, with primary antibodies (**Table S1**). Incubation with secondary antibodies and nuclei counterstaining were performed at room temperature (RT) followed by mounting in Mowiol fluorescent medium. Fluorescence images were acquired with the Cell Observer SD spinning disk confocal microscope (Carl Zeiss, Germany) and the number of single- and double-positive cells calculated using Zeiss ZEN 2.1 software (8-10 sections per animal and 4-5 animals per group). Details regarding tissue processing, immunostainings, image acquisition and stereology are further described in supplementary information.

### 2.4. Biochemical and molecular analyses

#### 2.4.1. Corticosterone measurement

A commercially available corticosterone ELISA kit (Abcam, UK) was used to quantify serum corticosterone (n=8-10 per group) according to manufacturer instructions, as detailed in supplementary information.

#### 2.4.2. Gene Expression

Total RNA was isolated from dissected DGs (n=6 per group) using RiboZol™ reagent method, according to the manufacturer’s instructions (VWR Life Science, USA). A total of 400 ng RNA was reversibly transcribed into cDNA using Xpert cDNA Synthesis Mastermix kit (GRiSP) under manufacturer’s instructions. β-actin was used as an endogenous control to normalize expression levels of different genes described in **Table S3** and relative mRNA concentrations was measured using ΔΔCt comparative method.

#### 2.4.3. Protein Levels

Western blotting analysis was used to assess CB2R protein levels from DG tissue (n=4 per group) (see supplementary information for procedure details).

### 2.5. Electrophysiological recordings

After transcardial perfusion with PBS, extracted hippocampal brain slices were prepared and transferred onto recovery chambers filled with oxygenated artificial cerebrospinal fluid (aCSF). Evoked field excitatory postsynaptic potentials (fEPSPs) were recorded in the outer molecular layer of the DG, similar to previous studies [55], as described in supplementary information.

### 2.6. Data analysis and statistics

Group sample sizes were based on numbers using uCMS models [46] and normality was assessed using the Shapiro-Wilk statistical test. Data were analysed through two-tailed unpaired and paired student’s t-tests, one-way, two-way or three-way analysis of variance (ANOVA), with Dunnett’s correction for multiple comparisons when appropriate (unless stated otherwise). Results are expressed as mean ± standard error of mean (SEM) and statistical significance was set when *p*<0.05. Statistical analysis was performed using Prism v.8.4.2 (GraphPad Software, US). For brevity, test details are described in figure captions, supplementary information and **Table S4** (*p*<0.05 values highlighted in bold and italics).

## 3. Results

### 3.1. CB2R inhibition in combination with PE counteracts chronic stress-induced behavioural changes

First, the best dosage scheme for CB2R modulation was evaluated utilizing the CB2R agonist HU308 and the CB2R inverse agonist AM630 (**Fig. S1a**). In physiological conditions, HU308 administration at different doses (0.5 and 5 mg/kg) did not exert any significant effect in anxiety- and depressive-like behaviours or cognitive performance (**Fig. S1b,c,f**). When assessing neurogenesis through BrdU injections four weeks before sacrifice, HU308 administration was found to increase the number of cells expressing the immature neuronal marker DCX (**Fig. S1h**) and the number of BrdU+/DCX+ cells (**Fig. S1h,j**), suggesting an effect on early neuronal differentiation. Conversely, AM630 administration at different doses (0.5 and 5 mg/kg) decreased time spent in open arms in the EPM, increased immobility in the FST, and compromised novel object recognition in the NOR test, indicating induction of anxiety- and depressive-like behaviours and impaired cognitive performance (**Fig. S1d,e,g**). However, AM630 administration did not affect the number of DCX+ or BrdU+/DCX+ cells in the DG, indicating a lack of effect on AHN at these doses (**Fig. S1i,j**). Considering these results and the fact that most previous studies report effects using low doses of CB2R ligands [25, 28, 45], the following experiments with the uCMS were performed using the lowest dose for each CB2R ligand (0.5 mg/kg).

To test our hypothesis, an uCMS protocol [48] was executed for 8 weeks followed by the combined strategy treatment and behavioural assessment for depressive-like behaviour, anxiety and cognition (**Fig. 1a**; further details in **Fig. S2a,b**). Briefly, uCMS exposure negatively impacted weight gain (**Fig. S2c**) and increased nadir corticosterone levels in the experimental conditions (**Fig. S2d,e**). Of note, uCMS enhanced paired-pulse inhibition (**Fig. S2f**) and promoted a tendency towards reduced CB2R levels (p=0.052) in the DG when comparing with CTRL conditions (**Fig. S2g**).

**Fig. 1.**
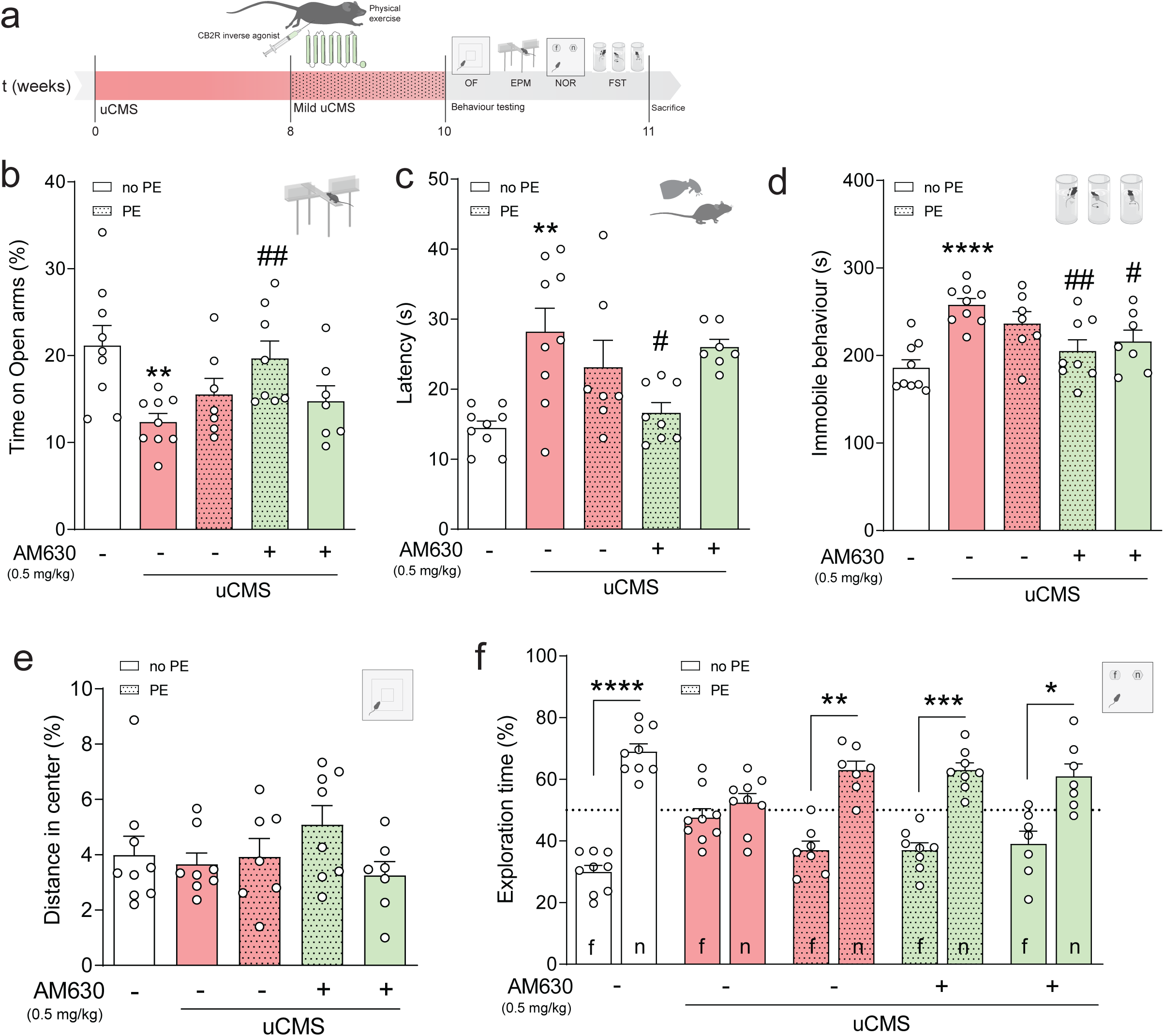
CB2R inhibition in combination with PE counteracts chronic stress-induced behavioural impairments. **a** Schematic representation of the experimental design. **b** uCMS induces anxiety-like behaviour (decrease in time spent in open arms during EPM) that is ameliorated by AM630 treatment in combination with PE. **c** Increased latency to groom in the SST evoked by uCMS returns to similar levels as CTRL with AM630 treatment in combination with PE. **d** uCMS-induced immobile behaviour in FST is decreased in animals subjected to AM630 treatment in combination with PE. **e** uCMS and subsequent treatments do not change the time in the centre zone of the OF test. **f** uCMS-induced cognitive impairments in NOR are ameliorated by PE alone or in combination with AM630 treatment. Data presented as mean ± SEM and circles individual data points (animals) [Student’s t-test: ****p<0.0001, ***p<0.001, **p<0.01, *p<0.05 vs CTRL; Two-way ANOVA: ^##^p<0.01; ^#^p<0.05 vs uCMS; further statistical details in **Table S4**]. uCMS, unpredictable chronic mild stress; AM630, CB2R inverse agonist; PE, physical exercise; no PE, no physical exercise.

As previously shown [56], chronic stress elicited an anxiety-like phenotype associated with a decreased time spent in the open arms of the EPM test (**Fig. 1b**). This effect was counteracted by AM630 administration in combination with PE to levels similar to non-stressed control (CTRL) animals (**Fig. 1b**). Contrarily, this effect was not observed with HU308 treatment (**Fig. S3b**). Mice exposed to uCMS also exhibited an increased latency to groom in the SST, indicating an anhedonic-like behaviour, an effect that was ameliorated by AM630 administration in combination with PE (**Fig. 1c**). HU308 administration, however, did not improve self-care behaviour in chronically stressed mice (**Fig. S3c**). Concomitantly, uCMS exposure greatly disrupted the ability to cope with an inescapable stressor in the FST (**Fig. 1d**). This stress-evoked disruption was counteracted by treatment with either AM630 alone or, more markedly, in combination with PE (**Fig. 1d**), but not by HU308 administration (**Fig. S3d**). No changes in the distance travelled in the centre of the arena were observed in the OF test, another anxiety-related paradigm (**Fig. 1e**). Similar results were obtained when assessing the influence of HU308 administration on OF performance (**Fig. S3e**). Finally, when looking at memory function, chronically stressed mice displayed significant deficits in the NOR test, as denoted by a decreased preference to explore the novel object (**Fig. 1f**). Treatment with AM630, PE, or a combination of both prevented these stress-induced cognitive changes (**Fig. 1f**) whilst HU308 administration or in combination with PE did not (**Fig. S3f**).

Collectively, these observations show that CB2R inverse agonist treatment or PE alone do not ameliorate most emotional behavioural deficits induced by uCMS exposure. Importantly, AM630 administration in combination with PE prevented the deficits in emotional behaviour, suggesting that CB2R inhibition is instrumental for PE to fully exert its beneficial actions in counteracting the effects of chronic stress on mood regulation.

### 3.2. CB2R inhibition in combination with PE counteracts the deficits in AHN induced by chronic stress

Given the role of AHN in emotional and memory functions [4], we next assessed whether AM630 administration in combination with PE induced changes in AHN in chronically stressed mice. Of note, estimated DG volumes did not vary between tested experimental conditions (**Table S2**). Stereological analysis revealed that uCMS exposure significantly decreased the number of proliferating cells (Ki67+), and this effect was counteracted by AM630 treatment in combination with PE (**Fig. 2a,d**). The reduction in the number of proliferating cells induced by uCMS exposure was particularly marked in the vHip (**Fig. 2a3**), reinforcing previous evidence indicating that chronic stress preferentially targets the ventral AHN [57, 58].

**Fig. 2.**
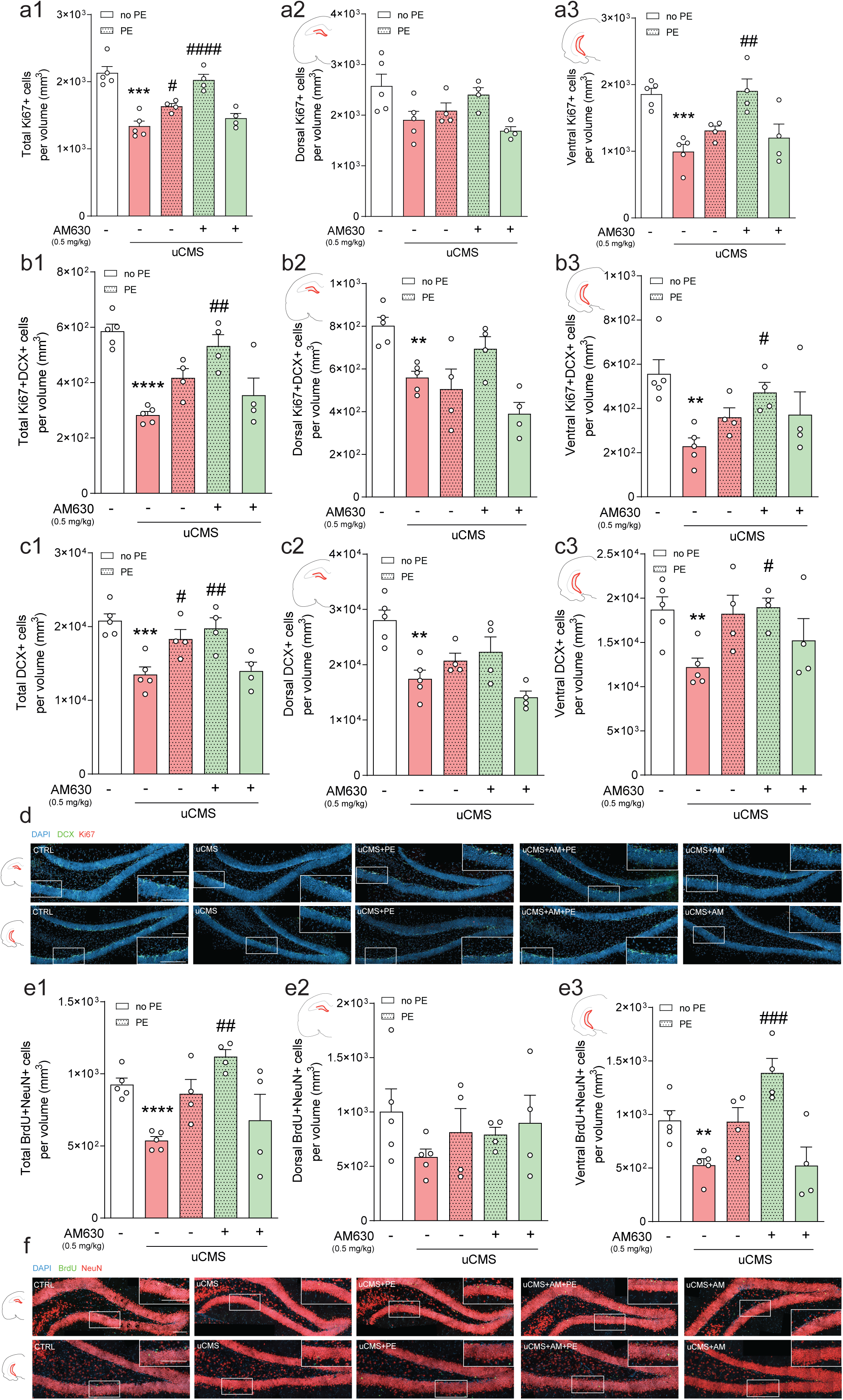
CB2R inhibition in combination with PE ameliorates chronic stress-induced negative impacts in AHN. **a** Chronic stress decreases the total number of proliferating (Ki67+) cells, particularly evident in the vHip, and AM630 treatment in combination with PE significantly rescues this deficit. Cell counts of Ki67+ cells per total (a1), dorsal (a2) and ventral (a3) estimated volumes are depicted. **b** Chronic stress decreases the number of proliferating neuroblasts (Ki67+DCX+), and AM630 treatment in combination with PE counteracts the stress-induced detrimental effect. Cell counts of Ki67+DCX+ cells per total (b1), dorsal (b2) and ventral (b3) estimated volumes are depicted. **c** The pool of immature neurons (DCX+) is diminished by uCMS, an effect countered by PE alone or in combination with AM630 treatment in combination and particularly evident in the vHip. Cell counts of DCX+ cells per total (c1), dorsal (c2) and ventral (c3) estimated volumes are depicted. **d** Representative images of coronal sections of dHip and vHip of all tested conditions, depicting cells stained for DCX (green), Ki67 (red) and DAPI (blue) with magnified boxed areas showing the detailed expression of individual markers in each condition; scale bars = 50 µm. **e** Chronic stress significantly impairs the survival of newborn neurons (BrdU+NeuN+), an effect prevented by AM630 treatment in combination with PE, particularly evident in the vHip. Cell counts of BrdU+NeuN+ cells per total (e1), dorsal (e2) and ventral (e3) estimated volumes are depicted. **f** Representative images of coronal sections of dHip and vHip of all tested conditions, depicting cells stained for BrdU (green), NeuN (red) and DAPI (blue) with magnified boxed areas showing the detailed expression of individual markers in each condition; scale bars = 50 µm. Data presented as mean ± SEM and circles individual data points (animals) [Student’s t-test: \*\*\*\**p*<0.0001, \*\*\**p*<0.001, \*\**p*<0.01, \**p*<0.05 vs CTRL; Two-way ANOVA: ^####^*p*<0.0001, ^###^*p*<0.001, ^##^*p*<0.01, ^#^*p*<0.05 vs uCMS; further statistical details in **Table S4**]. uCMS, unpredictable chronic mild stress; AM630, CB2R inverse agonist; PE, physical exercise.

We observed a marked decrease in the number of proliferating neuroblasts (Ki67+DCX+) following uCMS exposure (**Fig. 2b,d**). This effect was evident in both dHip and vHip, although more pronounced in the vHip (**Fig. 2b2,b3**). Remarkably, the negative effects of uCMS upon proliferating neuroblasts were prevented by AM630 treatment in combination with PE (**Fig. 2b1**), so that treated animals presented levels similar to those of CTRL animals.

Similarly, the number of DCX+ cells was significantly affected by uCMS exposure (**Fig. 2c,d**). Interestingly, this effect was counteracted by AM630 treatment in combination with PE, but not AM630 alone (**Fig. 2c1**), and again, the beneficial effects exerted by the combined treatment were particularly evident in the vHip (**Fig. 2c3**).

To evaluate the survival of newborn neurons, animals were injected with BrdU four weeks prior to sacrifice to evaluate the survival of newborn neurons (**Fig. S2a**) and cell fate was determined by co-localization of BrdU+ cells with NeuN (marker of mature neurons). We observed that uCMS exposure greatly affected the survival of newborn neurons in the DG, reducing the number of BrdU+NeuN+ cells (**Fig. 2e,f**), an effect completely counteracted by the combination of AM630 treatment with PE (**Fig. 2e,f**). These stress-evoked changes and concomitant proneurogenic actions of our combined strategy were particularly marked in the vHip (**Fig. 2e3**).

Overall, these observations suggest that chronic stress-induced behavioural deficits are accompanied by perturbations in the proliferation and survival of adult-born hippocampal neurons, particularly evident in the vHip. Importantly, although PE alone partially ameliorates some of these negative changes, only the CB2R inverse agonist treatment in combination with PE was able to fully counteract AHN deficits at all AHN stages (to similar levels as CTRL animals), strengthening the argument that CB2Rs are crucial to regulate the positive effects mediated by PE following chronic stress.

### 3.3. CB2R inhibition in combination with PE rescues chronic stress-induced changes in AHN in a subregional manner

To further clarify the impact of AM630 administration in combination with PE on the recovery of neurogenic disturbances elicited by chronic stress, we dissected the subregional distribution of the different cell populations related to AHN, considering the functional differentiation of adult-born neurons along longitudinal and transverse axes [56, 57]. First, we observed that Ki67+ cells distributed along the supra- and infrapyramidal blades in a ∼3:2 proportion, with uCMS exposure disrupting this regional distribution (**Fig. 3a**, **Table 1**). Specifically, uCMS promoted an overall change in the distribution of Ki67+ cells from the suprapyramidal SGZ to the corresponding granular cell layer (GCL) in both dHip and vHip. This effect was attenuated by PE alone or AM630 treatment in combination with PE (**Fig. 3a**, **Table 1**).

**Fig. 3.**
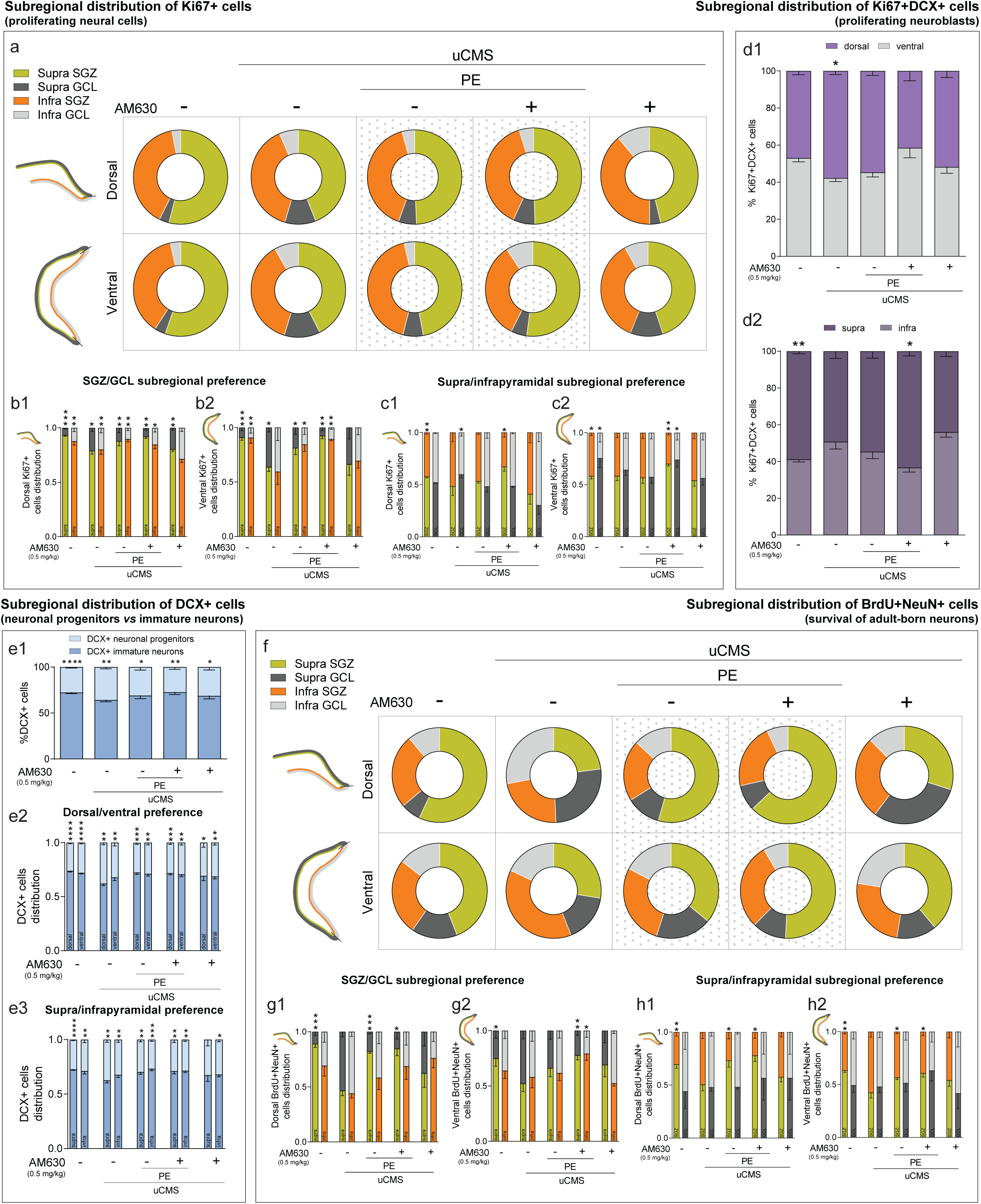
Chronic stress-induced disruptions in DG regional distribution of proliferating cells and newborn neurons are counteracted by CB2R inhibition in combination with PE. **a** Chronic stress disrupts dorsal and ventral regional distribution of proliferating (Ki67+) cells in DG subregions and AM630 treatment in combination with PE restores regional disturbances. **b** Quantitative analysis of the dorsal (b1) and ventral (b2) SGZ/GCL subregional preference for the location of proliferating cells. **c** Quantitative analysis of the dorsal (c1) and ventral (c2) supra/infrapyramidal subregional preference for the location of proliferating cells. **d** Chronic stress disrupts dorsal/ventral (d1) and supra/infra (d2) regional distribution of proliferating neuroblasts (Ki67+DCX+), that is rescued by AM630 treatment in combination with PE. **e** Chronic stress alters the balance between progenitor and immature DCX+ cells, which is restored by AM630 treatment in combination with PE (e1); quantitative analysis of the dorsal/ventral (e2) and supra/infra (e3) subregional preference for the location of DCX+ cells. **f** Chronic stress disrupts dorsal and ventral regional distribution of newly born neurons (BrdU+NeuN+) in different DG subregions; AM630 treatment in combination with PE restores these regional disturbances. **g** Quantitative analysis of the dorsal (g1) and ventral (g2) SGZ/GCL subregional preference for the location of newly born neurons. **h** Quantitative analysis of the dorsal (h1) and ventral (h2) supra/infrapyramidal subregional preference for the location of newly born neurons. Data presented as mean ± SEM (except for donut plots where only mean is presented) [Student’s t-test: \*\*\*\**p*<0.0001. \*\*\**p*<0.001, \*\**p*<0.01, \**p*<0.05; further statistical details in **Table S4**]. Supra, suprapyramidal blade; Infra, infrapyramidal blade; SGZ, subgranular zone; GCL, granule cell layer; uCMS, unpredictable chronic mild stress; AM630, CB2R inverse agonist; PE, physical exercise.

**Table 1.**
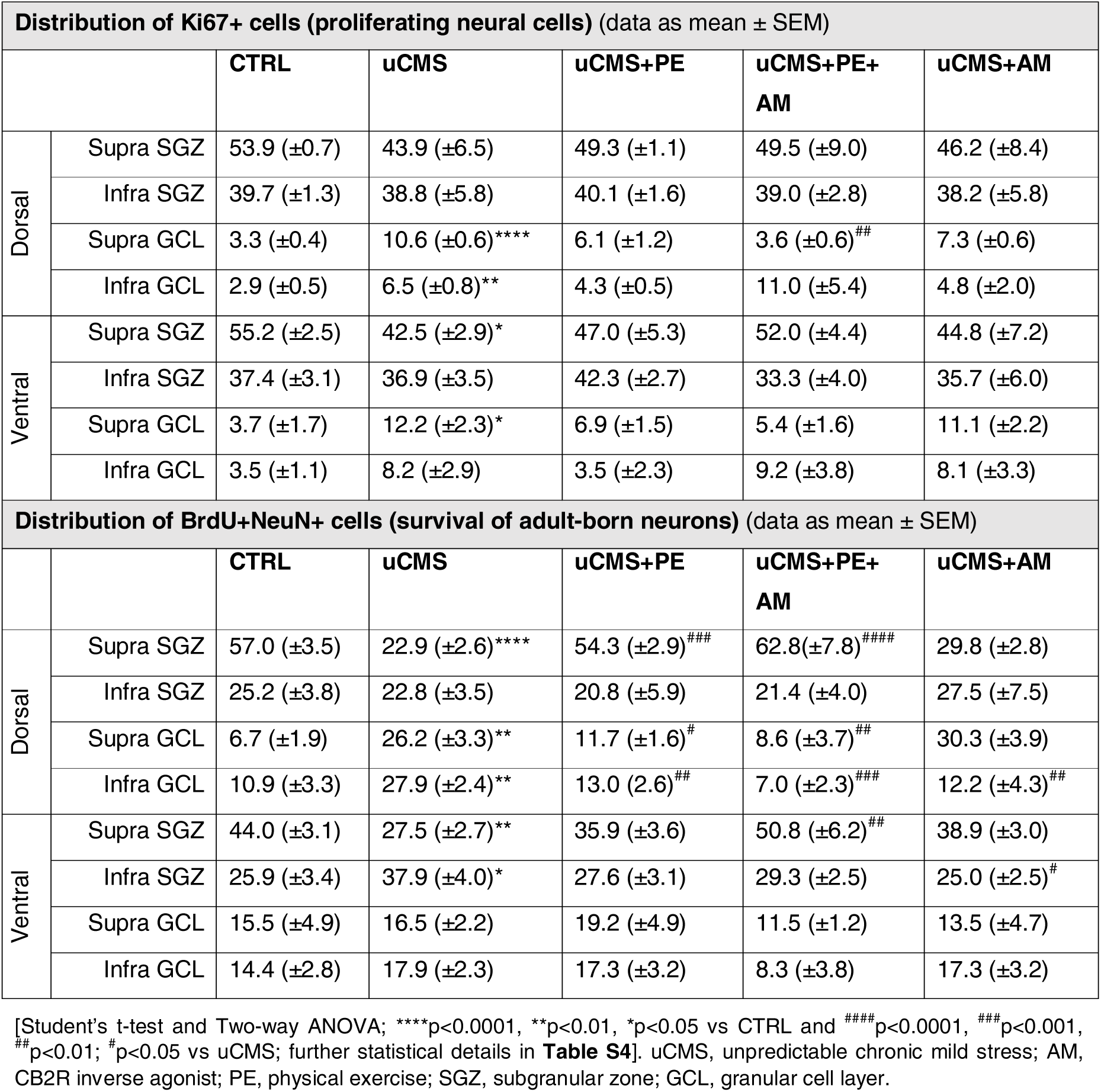
Total regional distribution of neural populations in DG subregions in tested experimental conditions

| Distribution of Ki67+ cells (proliferating neural cells) (data as mean ± SEM) |  |  |  |  |  |  |
| --- | --- | --- | --- | --- | --- | --- |
|  |  | CTRL | uCMS | uCMS+PE | uCMS+PE+AM | uCMS+AM |
| Dorsal | Supra SGZ | 53.9 (±0.7) | 43.9 (±6.5) | 49.3 (±1.1) | 49.5 (±9.0) | 46.2 (±8.4) |
|  | Infra SGZ | 39.7 (±1.3) | 38.8 (±5.8) | 40.1 (±1.6) | 39.0 (±2.8) | 38.2 (±5.8) |
|  | Supra GCL | 3.3 (±0.4) | 10.6 (±0.6)**** | 6.1 (±1.2) | 3.6 (±0.6)## | 7.3 (±0.6) |
|  | Infra GCL | 2.9 (±0.5) | 6.5 (±0.8)** | 4.3 (±0.5) | 11.0 (±5.4) | 4.8 (±2.0) |
| Ventral | Supra SGZ | 55.2 (±2.5) | 42.5 (±2.9)* | 47.0 (±5.3) | 52.0 (±4.4) | 44.8 (±7.2) |
|  | Infra SGZ | 37.4 (±3.1) | 36.9 (±3.5) | 42.3 (±2.7) | 33.3 (±4.0) | 35.7 (±6.0) |
|  | Supra GCL | 3.7 (±1.7) | 12.2 (±2.3)* | 6.9 (±1.5) | 5.4 (±1.6) | 11.1 (±2.2) |
|  | Infra GCL | 3.5 (±1.1) | 8.2 (±2.9) | 3.5 (±2.3) | 9.2 (±3.8) | 8.1 (±3.3) |
| Distribution of BrdU+NeuN+ cells (survival of adult-born neurons) (data as mean ± SEM) |  |  |  |  |  |  |
|  |  | CTRL | uCMS | uCMS+PE | uCMS+PE+AM | uCMS+AM |
| Dorsal | Supra SGZ | 57.0 (±3.5) | 22.9 (±2.6)**** | 54.3 (±2.9)### | 62.8(±7.8)#### | 29.8 (±2.8) |
|  | Infra SGZ | 25.2 (±3.8) | 22.8 (±3.5) | 20.8 (±5.9) | 21.4 (±4.0) | 27.5 (±7.5) |
|  | Supra GCL | 6.7 (±1.9) | 26.2 (±3.3)** | 11.7 (±1.6)# | 8.6 (±3.7)## | 30.3 (±3.9) |
|  | Infra GCL | 10.9 (±3.3) | 27.9 (±2.4)** | 13.0 (2.6)## | 7.0 (±2.3)### | 12.2 (±4.3)## |
| Ventral | Supra SGZ | 44.0 (±3.1) | 27.5 (±2.7)** | 35.9 (±3.6) | 50.8 (±6.2)## | 38.9 (±3.0) |
|  | Infra SGZ | 25.9 (±3.4) | 37.9 (±4.0)* | 27.6 (±3.1) | 29.3 (±2.5) | 25.0 (±2.5)# |
|  | Supra GCL | 15.5 (±4.9) | 16.5 (±2.2) | 19.2 (±4.9) | 11.5 (±1.2) | 13.5 (±4.7) |
|  | Infra GCL | 14.4 (±2.8) | 17.9 (±2.3) | 17.3 (±3.2) | 8.3 (±3.8) | 17.3 (±3.2) |

When looking at the subregional distribution of proliferating cells along DG transverse axis, we observed that the majority of Ki67+ cells was located at the SGZ (∼95%) of both supra- and infrapyramidal blades in the dHip, as previously reported [56]. uCMS exposure followed by AM630 treatment in combination with PE affected this distribution (**Fig. 3b1**). Interestingly, we found that uCMS exposure critically affected the subregional positioning of proliferating cells in the SGZ and GCL subregions in both supra- and infrapyramidal blades of the vHip, an effect countered by PE alone or in combination with AM630 treatment (**Fig. 3b2**). We also observed differences between the supra- and infrapyramidal blades of both dHip (**Fig. 3c1**) and vHip (**Fig. 3c2**) when looking at SGZ and GCL subregions separately. Specifically, while CTRL animals displayed slightly higher levels of Ki67+ cells in the SGZ and GCL of the suprapyramidal layer, in both dHip and vHip, uCMS exposure changed this ratio to similar values in both layers (**Fig. 3c**). Strikingly, AM630 treatment in combination with PE restored the distribution of Ki67+ cells, an effect particularly evident in the vHip (**Fig. 3c2**).

Next, we quantified proliferating neuroblasts (Ki67+DCX+ cells) across the septotemporal axis. We observed a similar distribution of Ki67+DCX+ cells in the dHip (∼47%) and vHip (∼53%) in CTRL animals. However, uCMS exposure changed this proportion (∼58% dHip; ∼42% vHip) while the numbers of Ki67+DCX+ cells were not significantly different from CTRL after AM630 treatment in combination with PE (**Fig. 3d1**). Ki67+DCX+ cells were preferentially located in the suprapyramidal blade along the transverse axis (∼60%) in CTRL conditions. Chronic stress altered this distribution, favouring the presence of more Ki67+DCX+ cells in the infrapyramidal blade, and AM630 treatment in combination with PE prevented this effect (**Fig. 3d2**).

To quantify the effects of uCMS exposure and our treatments on neuronal progenitors and immature neurons in the DG, we counted: 1) DCX+ cells located in the SGZ that exhibited no processes or short processes parallel to the GCL (neuronal progenitors), and 2) DCX+ cells in the two internal thirds of the GCL that exhibited a branched dendritic extension radially crossing the GCL (immature neurons), as previously described [59] (**Fig. 3e**). We found that, ∼25% of DCX+ cells corresponded to neuronal progenitors and ∼75% to immature neurons in CTRL conditions. uCMS exposure increased the proportion of neuronal progenitors to ∼34%. The effect of uCMS was reverted by PE alone or in combination with AM630 treatment (**Fig. 3e1**). Interestingly, we observed that the aforementioned ratios were maintained in both dHip (∼26% of neuronal progenitors and ∼74% of immature neurons) and vHip (∼28% of neuronal progenitors and ∼72% of immature neurons) along the septotemporal axis, with uCMS exposure drastically increasing the number of neuronal progenitors, particularly in the dHip (∼38% of neuronal progenitors and ∼62% of immature neurons) compared to CTRL (**Fig. 3e2**). Similarly, the same effects were observed along the transverse axis, with the 1:3 proportion of DCX+ progenitor to immature cells in supra- and infrapyramidal blades being disturbed by uCMS exposure. Specifically, the suprapyramidal ratio was affected by uCMS exposure, which slightly increased the number of progenitor cells (∼37% of neuronal progenitors) compared to CTRL levels (∼27% of neuronal progenitors). This effect was rescued by PE alone or in combination with AM630 treatment (**Fig. 3e3**).

Finally, we studied the survival of newborn neurons by quantifying the numbers of BrdU+NeuN+ cells in the DG. We observed that BrdU+NeuN+ cells were distributed along the supra- and infrapyramidal blades in a ∼3:2 proportion, with uCMS exposure greatly impacting this regional distribution. Specifically, uCMS altered the numbers of BrdU+NeuN+ cells in supra- and infrapyramidal blades and their location within the GCL (**Fig. 3f**, **Table 1**). Importantly, the numbers and location of BrdU+NeuN+ cells were similar to CTRL conditions in uCMS mice treated with AM630 in combination with PE (**Fig. 3f**, **Table 1**). A closer examination of the distribution of BrdU+NeuN+ cells along the DG transverse axis revealed that uCMS exposure disrupted dHip SGZ/GCL subregional distribution of BrdU+NeuN+ cells (CTRL: ∼95% in supraSGZ and ∼70% in infraSGZ vs uCMS: ∼50% in supraSGZ and ∼45% in infraSGZ). This effect was reverted by PE alone or in combination with AM630 treatment (**Fig. 3g1**). Similarly, we found that uCMS critically affected the subregional distribution of BrdU+NeuN+ cells in vHip SGZ and GCL subregions, an effect particularly evident in the suprapyramidal blade and countered by AM630 treatment in combination with PE (**Fig. 3g2**). When looking at SGZ and GCL subregions separately, we observed that uCMS exposure disrupted the distribution of BrdU+NeuN+ cells in supra- and infrapyramidal blades. This effect, particularly relevant in the SGZ arrangement, was attenuated in the dHip (**Fig. 3h1**) and the vHip (**Fig. 3h2**) by PE alone or in combination with AM630 treatment.

Altogether, these results reveal that uCMS exposure disturbs the regional distribution of AHN-derived cell populations, promoting the relocation of proliferating cells (Ki67+ cells) and newborn neurons (Ki67+DCX+ and BrdU+NeuN+ cells) and differentially affecting the distribution of these cell populations. Importantly, although PE alone was able to revert some of these features, co-treatment with AM630 boosted this effect, normalizing proportions to levels similar to those of CTRL animals. This suggests that CB2R inverse agonist treatment in combination with PE restores the regional balance of AHN-derived cell populations, which was impaired by chronic stress.

### 3.4. Neuroinflammation-related changes evoked by chronic stress are prevented by CB2R inhibition in combination with PE

The effects in the different behavioural domains together with changes in AHN that we described in the previous sections, prompted us to study the overall expression of microglia, astrocytes and myelin in the granular layer of the DG, as they may act as supportive cell types [6]. Moreover, given the anti-inflammatory properties of both PE and CB2Rs [25, 37] the expression of different neuroinflammatory markers was evaluated.

First, we looked at changes promoted by chronic stress and our combined strategy treatment in microglia expression as indicated by ionized calcium binding adaptor molecule 1 (Iba1), myelin expression as indicated by myelin basic protein (MBP) and astrocytic expression as indicated by astrocytic glial fibrillary acidic protein (GFAP) (**Fig. 4a-d**). Analysis of DG granular layer revealed that uCMS induced an increase in Iba1 immunoreactivity, suggestive of enhanced microgliosis. Remarkably, PE alone or in combination with AM630 treatment reverted this effect (**Fig. 4a,d**). Additionally, the combination of HU308 and PE reduced Iba1 immunoreactivity, but HU308 alone did not exert any significant effects on Iba1 immunoreactivity (**Fig. S4a**). uCMS also significantly reduced MBP immunoreactivity in the granular zone. This effect was prevented by AM630 alone or in combination with PE (**Fig. 4b,d**), but not by HU308 treatment (**Fig. S4b**). No alterations in GFAP immunoreactivity were observed between groups (**Fig. 4c,d** and **Fig. S4c**).

**Fig. 4.**
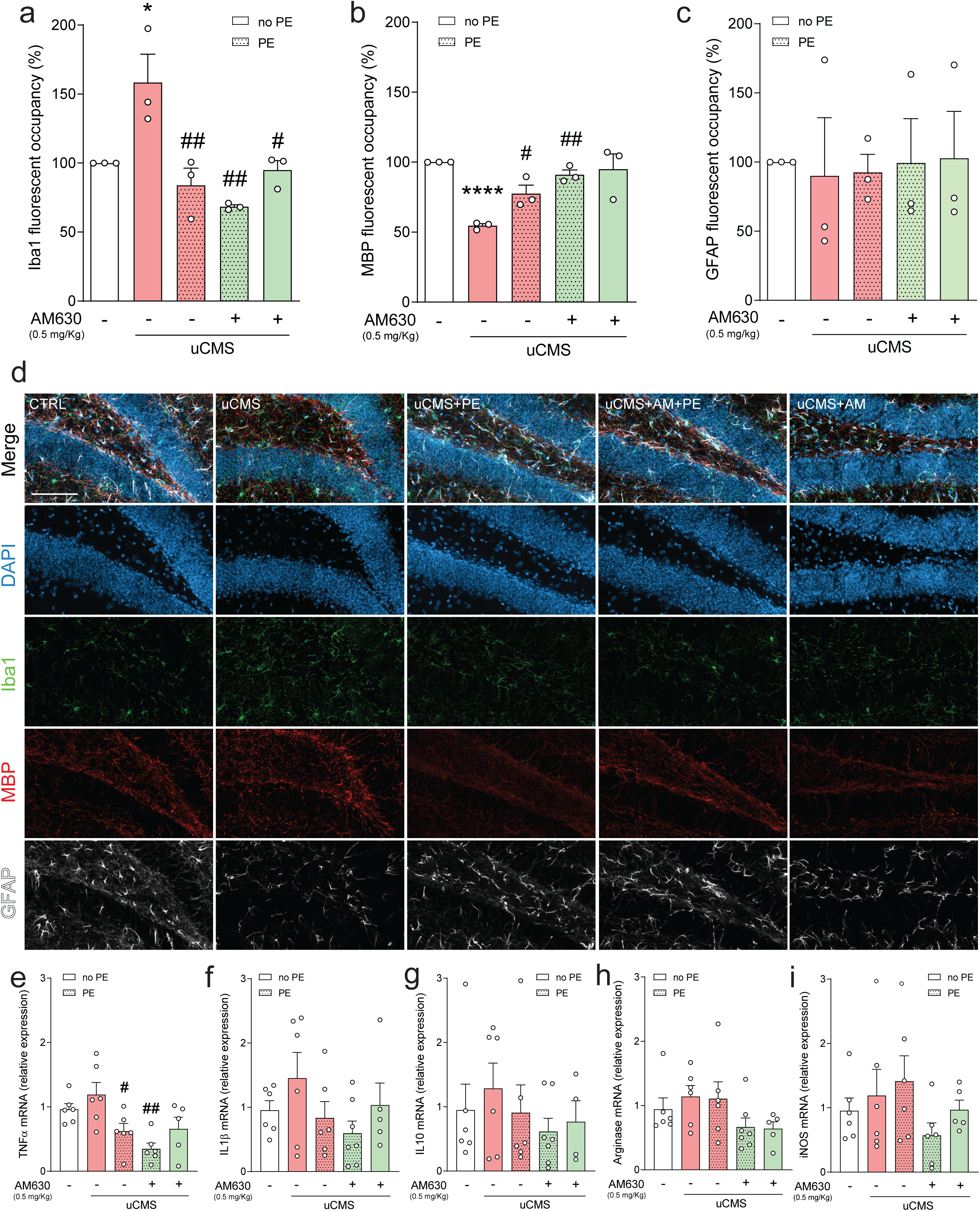
CB2R inhibition in combination with PE ameliorates chronic stress-induced neuroinflammatory changes. **a** Chronic stress increases Iba1 granular expression, an effect prevented by PE alone, AM630 treatment or a combination of both. **b** Chronic stress-induced decrease in MBP granular expression is rescued by PE alone or in combination with AM630 treatment. **c** GFAP granular expression remains unchanged after uCMS or PE in combination with AM630 treatment. **d** Representative images of coronal sections of hippocampal DG of all tested conditions, stained for DAPI (blue), Iba1 (green), MBP (red) and GFAP (white) with merged and individual channels; scale bar = 50 µm. The expression of neuroinflammation-related markers **e** TNFα, **f** IL1β, **g** IL10, **h** arginase and **i** iNOS is differentially affected by uCMS and PE in combination with AM630 treatment. Data presented as mean ± SEM and circles individual data points (animals) [Student’s t-test: \*\*\*\**p*<0.0001, \*\**p*<0.01, \**p*<0.05 vs CTRL; Two-way ANOVA: ^##^*p*<0.01, ^#^*p*<0.05 vs uCMS; further statistical details in **Table S4**]. uCMS, unpredictable chronic mild stress; AM630, CB2R inverse agonist; PE, physical exercise.

The changes in immunoreactivity of cell-type specific markers were accompanied by alterations in different neuroinflammatory markers in the DG. Despite not observing significant differences in the relative expression of first-line pro-inflammatory cytokines in the DG such as tumour necrosis factor α (TNFα), interleukin 1β (IL-1β) and anti-inflammatory interleukin 10 (IL-10) after uCMS exposure, TNFα levels were reduced by PE treatment and even more by its combination with AM630 treatment (**Fig. 4e-g**), likely indicating a role for this combined strategy in controlling early neuroinflammation. Although uCMS had no detectable effect on the expression of enzymes related to microglial reactivity and neuroinflammation, arginase 1 (arginase) and inducible nitric oxide synthase (iNOS), AM630 treatment in combination with PE reduced their expression when compared to uCMS (**Fig. 4h,i**).

Overall, these observations indicate that, although the molecular machinery involved in neuroinflammation remains fairly unchanged, chronic stress affects the density of the glial cellular milieu in the DG. The neuroinflammatory load elicited by chronic stress upon microglial and astrocytic populations and changes in myelination were reduced by treatment with CB2R inverse agonist in combination with PE, suggesting that this strategy may be useful to counteract the stress-induced neuroinflammatory response in the DG.

## Discussion

In this study we uncover the role of CB2Rs in regulating PE-mediated actions and countering chronic stress behavioural and plasticity-induced defects. Taken together, our data show that inhibiting CB2Rs in combination with PE can restore the reduction in AHN and counteract the emotional and cognitive deficits impinged by chronic stress. Furthermore, we show that adjuvant CB2R inverse agonist treatment greatly boosts the positive effects of PE, resulting in behavioural recovery of emotional and cognitive dimensions (**Fig. 5a, a’**), increased NSC proliferation, neuronal differentiation and survival (**Fig. 5b, b’**) as well as decreased glial reactivity, myelination and neuroinflammation (**Fig. 5c, c’**). These observations suggest a pivotal participation of CB2Rs in PE-elicited recovery from chronic stress. Interestingly, a constant finding herein is that CB2R activation alone or in combination with PE yields no positive actions under chronic stress, in comparison with the beneficial effects of CB2R inhibition. Since in physiological conditions CB2R inhibition has a negative impact, our findings highlight a dual role of CB2Rs under physiological *versus* pathological conditions.

**Fig. 5.**
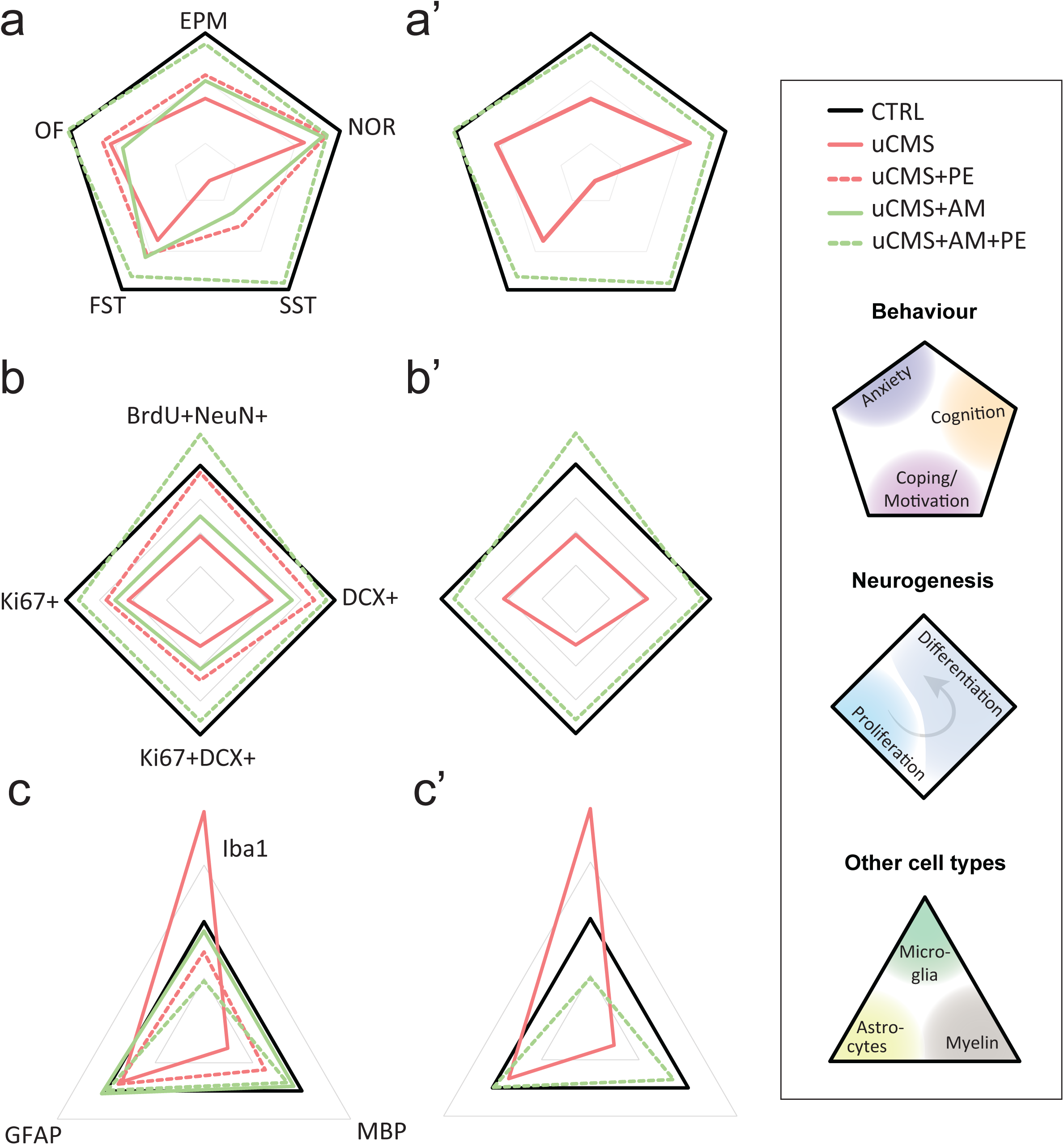
CB2R inhibition in combination with PE promotes robust antidepressant and proneurogenic effects after chronic stress. **a** Changes in different behavioural dimensions (anxiety-, depression- and cognitive-related behaviours) in all conditions (a). Impairments caused by uCMS are almost fully reverted by AM630 treatment in combination with PE (a’). **b** AHN-related changes in all conditions (b). Disruptions induced by uCMS are prevented by AM630 treatment in combination with PE (b’). **c** Changes in microglial, myelin and astrocytic granular expression in all conditions (c). Chronic stress-induced neuroinflammation-related imbalances are ameliorated by AM630 treatment in combination with PE (c’). Data presented as radar plots with absolute mean values, CTRL (black bold line) set to 100% and relative axis scales varying in between plots: centre points starting at 10%, 20%, 40% and distance between bounds (light grey lines) of 30%, 20% and 30% in a, b and c plots, respectively.

Although the beneficial effects of PE have been long known [60], the molecular mechanisms underlying this response have remained unclear. As recently proposed [61], understanding the molecular and cellular effects of PE, particularly with regard to effects on AHN and emotional and cognitive functions, is crucial to develop new preventive or combating strategies that mimic or enhance the beneficial effects of PE. From a translational standpoint, the therapeutic potential of physical exercise is enormous for the treatment of a variety of brain disorders (e.g., addiction, anxiety, stroke, epilepsy), as previous studies have highlighted [49, 62]. Current literature suggests that PE-elicited effects are mediated by the ECS [38, 41, 42]. In the context of stress, this interaction likely occurs due to PE mobilizing the ECS to replenish energy stores and participate in PE-induced analgesic and mood-elevating effects, consistent with a role of the ECS in both activating and terminating the hypothalamic-pituitary-adrenal axis response to stress [63, 64].

Recent studies started tackling this interaction in the context of NSC regulation and stress-related disorders, with only a few drawing correlations in clinical settings [40, 42, 43, 65]. In fact, this interplay has been showed to rely on the collaboration between ECS components and BDNF, one of the major neurotrophic factors upregulated by PE. For instance, endogenous cannabinoid signalling, specifically through cannabinoid type 1 receptors (CB1Rs), was shown to be required for the effects of voluntary PE in hippocampal NSCs [66]. In line with this, we have previously reported that CB2Rs are necessary for BDNF-mediated NSC proliferation and neuronal differentiation [44]. More broadly, others have found that cannabinoids prevent depressive-like behaviours, an effect that was accompanied by alterations in BDNF expression, in a rat model of posttraumatic stress disorder [67]. Recently, CB2R inhibition was found to dampen BDNF signalling in stressed animals, suggesting a close interaction between CB2Rs and BDNF in emotional response [45]. Interestingly, evidence suggests that exercise-mediated runner’s high (i.e., relaxing state of euphoria after PE) might occur due to the recruitment of peripheral CB1Rs and CB2Rs but, so far, no study has ever focused on whether CB2Rs participate and/or cooperate with PE in NSC regulation and stress response.

Our data now expands this field of research by showing that CB2Rs actively contribute to stress response elicited by PE. Specifically, we show that CB2R inhibition, but not its activation, is essential for PE-mediated antidepressant, anxiolytic and pro-cognitive actions after chronic stress. We observed that, in physiological conditions, CB2R activation had proneurogenic effects while CB2R inhibition triggered depressive-like behaviour and cognitive impairments, which is in accordance with existing literature [68]. However, and importantly, we observed that CB2R inhibition in combination with PE counteracted chronic stress-induced behavioural impairments, especially in the emotional domain. While chronic stress promoted anxiety-like, anhedonic-like behaviours and impaired coping features, CB2R inhibition alone was able to partially recover deficits in adaptive behaviour (i.e., in the FST test) but only in combination with PE it could fully ameliorate other behavioural dimensions. This is in agreement with previous publications showing that CB2R inhibition blocks the effects of chronic stress in behaviour tests related to emotional processing [26, 45], and we are now adding evidence that PE synergizes with this action. Regarding memory function, we observed that both PE alone or in combination with CB2R inhibition reverted chronic stress-induced cognitive impairments, which goes in line with previous studies showing that PE enhances learning and memory processes [60] and that CB2R is important for long-term memory consolidation [68]. Nevertheless, others have also shown that CB2R activation can exert beneficial effects regarding cognitive performance in a mouse model of Alzheimer’s disease [69].

Characterization of the neurogenic process in the hippocampal DG revealed that, in agreement with previous reports [56, 57, 70], chronic stress significantly impacts all stages of AHN – cell proliferation, neurogenesis and adult-born neuron survival. Our data goes in line with existing evidence indicating that the AHN deficits induced by chronic stress can affect both dorsal and ventral divisions of the hippocampal DG [58, 70], but preferentially targets the vHip [71, 72]. Although stress may preferentially decrease neurogenesis in the vHip, mood-improving interventions can stimulate neurogenesis in both hippocampal divisions [13], thus countering the effects of stress. Indeed, in our experimental setting, PE alone partially rescued some stress-evoked deficits in AHN, particularly regarding the number of immature neurons and survival of newborn neurons. This is in agreement with previous literature demonstrating that PE increases the number of immature neurons and adult-born neurons following stress [73, 74] but in conflict with evidence showing that PE can also boost cell proliferation [74, 75]. However, PE was able to fully rescue AHN deficits at all evaluated stages only in combination with CB2R inverse agonist AM630, suggesting that a reduction in CB2R constitutive activity is essential to boost the beneficial influence of PE in countering chronic stress. Growing evidence shows that CB2Rs are key regulators of the neurogenic process [16, 44], particularly in pathological contexts [25, 45], although CB2R knockout mice appear to display stable AHN [76], likely due to compensatory mechanisms. In our hands, CB2R inhibition alone was not able to revert any AHN deficits, further reinforcing that CB2R constitutive activity impinges on the actions of PE to shape adult hippocampal NSC rates of proliferation and differentiation. Further, these results suggest that the combined CB2R inhibition and PE treatment triggers a rearrangement in the DG neurogenic microenvironment, potentiating AHN in both dHip and vHip, which would explain the improvement in both dorsal and ventral hippocampal-dependent functions (i.e., cognitive and emotional dimensions).

Most studies focusing on the impact of chronic stress on AHN often disregard the structural segregation along both the hippocampus longitudinal and transverse axes or differences between the supra- and infra-pyramidal blades of the GCL, although newborn neurons localised there differentially modulate inputs to the DG [56, 72, 77]. This functional differentiation is extremely important because it translates into different behavioural responses related to distinct hippocampal functions. The dHip is preferentially involved in cognitive processes, with dorsal granule neurons participating in contextual memory, whereas the vHip is more implicated in emotional processing, with ventral granule neurons contributing to suppress innate anxiety [9, 13, 78]. The DG can be further subdivided into the suprapyramidal blade, located between CA1 and CA3 regions, and the infrapyramidal blade on the opposite side of the hilus [78], with each blade being further dissected into the SGZ and GCL subregions, likely reflecting different contributions to distinct hippocampal functions, such as the processing of spatial and contextual information [77, 79].

While aimed at disentangling the subregional contribution of each cellular population in AHN, we found chronic stress to disrupt subregional distributions of ongoing proliferating cells and newly-generated neurons, an effect that was more evident in the suprapyramidal blades of both dHip and vHip. Accordingly, when looking at immature neurons, we found that chronic stress induced a numerical shift between immature neurons and progenitor cells, from supra-to more infrapyramidal portions and from SGZ to more GCL regions. Importantly, the CB2R inhibition in combination with PE was able to abrogate these subregional changes and recover proliferating and survival patterns to similar levels as control conditions. These results go in line with previous studies showing that stress preferentially targets proliferation in the SGZ area [80] as well as the survival of adult-born neurons in the suprapyramidal blade [56]. Further studies are required to comprehensively understand the importance of these regional differences. As previously suggested [81], CB2R inhibition is likely boosting the beneficial effects of PE in preventing the stress-induced activation of DG mature granule neurons, which is known to critically shape AHN dynamics, likely through BDNF signalling [82].

Neuroinflammation plays a relevant role in depression, with both MDD patients and preclinical models of depression often exhibiting high levels of neuroinflammatory markers [37]. In our hands, we observe that uCMS exposure induces increased glial reactivity, recapitulating previous findings [83, 84]. We detected significant alterations associated with chronic stress in the expression of microglia and myelin, but not astrocytes, in DG granular regions. Taking into consideration the role of CB2Rs in regulating immunity and inflammation [25] and the role of PE as a strong inhibitor of inflammation [85], we confirmed the immunomodulatory role of PE in combination with CB2R inhibition, but not activation, in fighting neuroinflammation by reverting most changes triggered by chronic stress. These results go in line with previous evidence showing that inhibition of CB2Rs attenuates the inflammatory load triggered by challenging situations [86]. Nevertheless, others have also demonstrated that CB2R activation alone prevents stress-induced neuroinflammatory responses [25]. We posit that, in our experimental conditions, CB2R inhibition may be shielding the system against stress-targeted reactive microglia (which express high levels of CB2Rs [87]), thus facilitating PE actions in limiting the neuroinflammatory damage caused by chronic stress. Conversely, PE may be modulating stress-disrupted cellular immunity and CB2R inhibition further alleviating this immunoinflammatory status [88].

Although our results are associative, we suggest that the combined treatment of CB2R inhibition with PE accelerates the incorporation of newborn neurons into the hippocampal circuitry which, in turn, facilitates adaptive and resilient behaviours [89]. However, it is plausible that PE and CB2R modulation are interfering with the critical period that allows appropriate differentiation and survival of adult-born neurons by supplying adult NSCs with proneurogenic, pro-survival supportive factors (e.g., BDNF) that counteract the effects of stress.

Interestingly, AM630 was suggested to act as protean ligand of CB2Rs, which may be reducing the probability of spontaneous activation of the receptor by favouring an active receptor conformation of lower efficacy [90, 91]. This may reconcile some apparently controversial evidence showing beneficial effects of both CB2R activation and inhibition in stress-related conditions, since a protean ligand may act as agonist in systems that are quiescent (no constitutive activity) or as inverse agonist in constitutively active systems [91]. This possibility highlights that distinct stress levels may affect CB2R constitutive activity. Considering that the therapeutic relevance protean ligands may have in setting receptor activity to a constant level [91], one may speculate that, in our results, stress dysregulates CB2R constitutive activity and that AM630 may contribute to activity adjustments towards normal levels. Nevertheless, our data consistently demonstrates a cumulative effect of CB2R inhibition with PE, which dampens the negative consequences of stress, in most cases nullifying them.

In conclusion, our work shows that a strategy coupling CB2R inhibition in combination with PE may be a useful approach to counteract the effects of chronic stress in terms of behaviour and plasticity-related events in the hippocampus. Overall, our observations highlight a new multimodal approach for the treatment of stress-related pathologies. Future studies should focus on understanding whether the effects of this joint treatment expand to other brain regions and which intracellular mediators are behind the actions of this combined strategy. PE and CB2R effects may, indeed, be converging at multiple signalling pathways or, perhaps, have a common effector element. Altogether, given the lack of effective treatments and the significant public health impact of stress-related psychiatric disorders, in particular MDD, the promise of combining lifestyle interventions such as PE with pharmacological targeting of CB2Rs seems relevant and should be further explored.

## Acknowledgments

The authors are grateful to the Rodent, Histology and Comparative Pathology and Bioimaging facilities at Instituto de Medicina Molecular João Lobo Antunes (Lisbon, Portugal). This work was supported by ISN Career Development Grant, IBRO Early Career Award and FMUL/GAPIC project no. 20200008 granted to SX and by Fundação para a Ciência e a Tecnologia (FCT) IF/01227/2015 granted to SX, 2020.02855.CEECIND and 2022.02201.PTDC granted to LP and projects PTDC/MED-PAT/2582/2021, UIDB/04138/2020, UIDP/04138/2020 granted to AF as well as fellowships SFRH/BD/129710/2017 and COVID/BD/151880/2021 (RSR), PD/BD/141784/2018 (DML), PD/BD/150343/2019 (SLP), PD/BD/150341/2019 (JMM) and 2020.04492.BD (JBM). Authors would like to thank European Union’s Horizon 2020 research and innovation programs H2020-WIDESPREAD-05-2020-Twinning (EpiEpinet, no. 952455) and ERA-NET-NEURON (EJTC 2016) granted to AMS and CPF, respectively.

## Author contributions

RSR conceived the study, designed and performed the experiments, analysed results and wrote the paper; JBM, SHV, AB, SLP, JMM, DML, FFR performed the experiments and participated in result discussion; EL-C, PB provided help in the experiments; AF, AMS, LP, CPF, SX participated in result discussion and interpretation, provided funding support and corrected the paper; SX conceived and supervised the study, provided funding support and wrote the paper.

## Conflict of Interest

The authors declare no potential conflicts of interest.

## Data Availability Statement

The data supporting the findings are available from the corresponding author upon reasonable request.

---

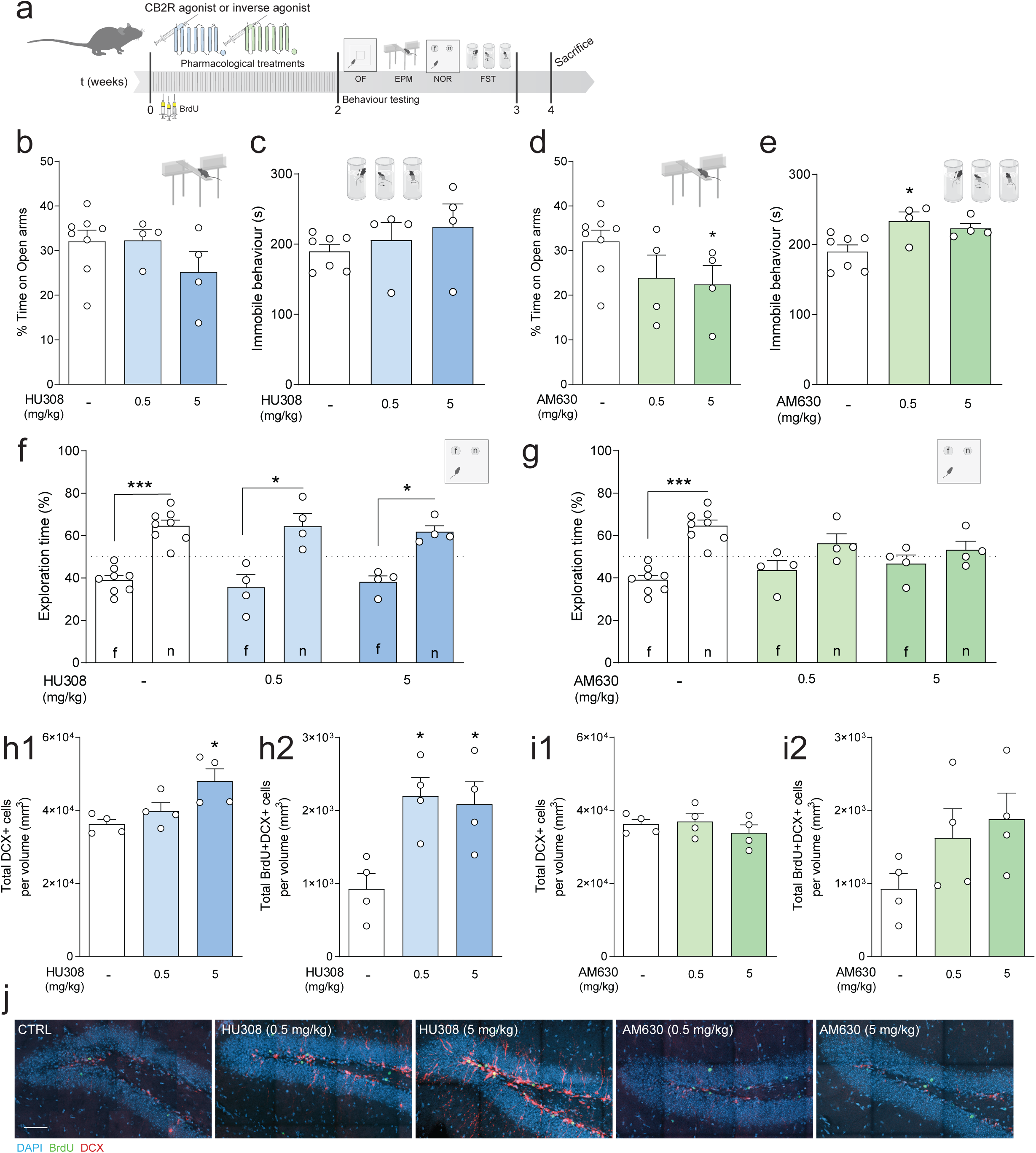

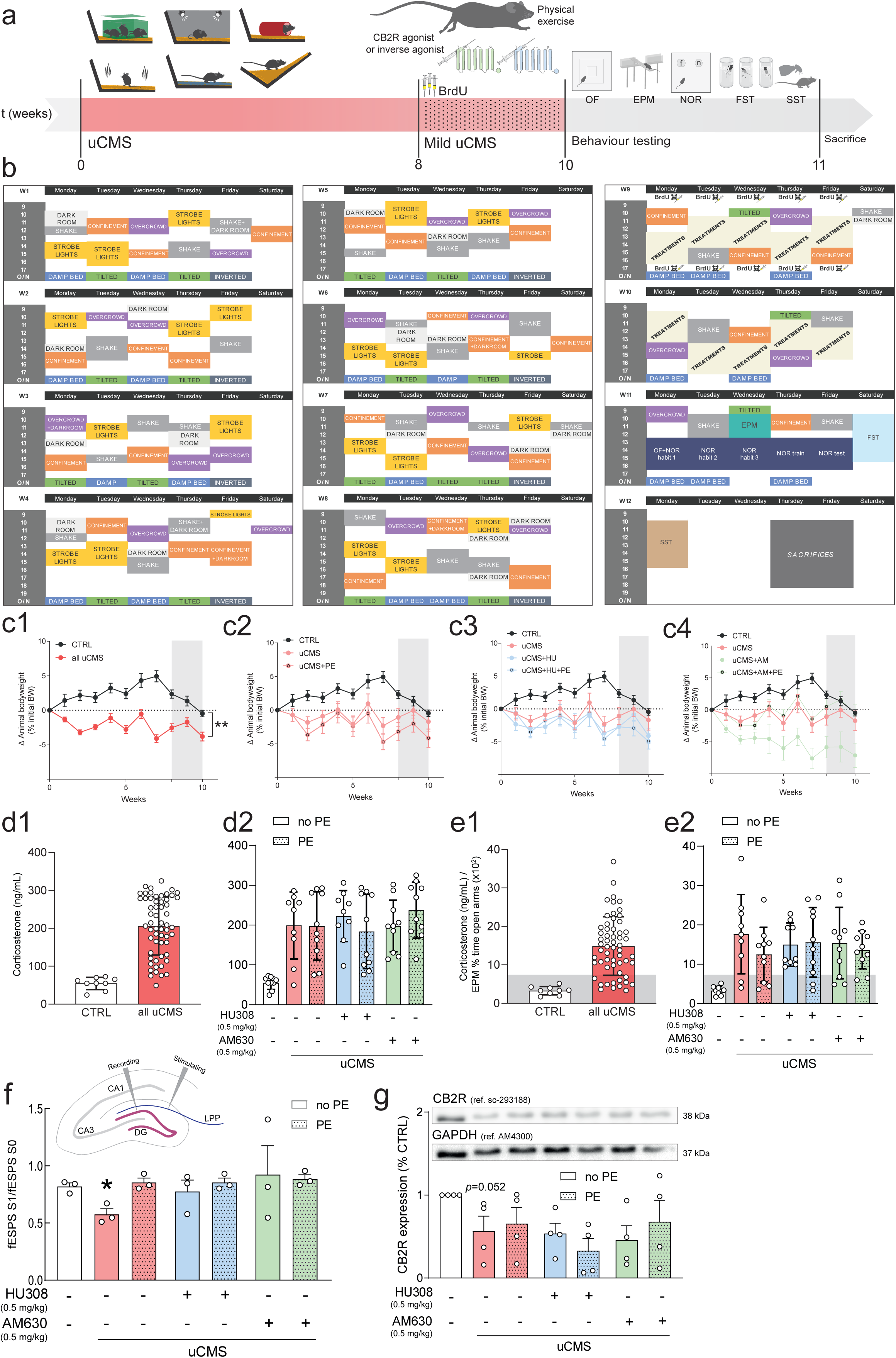

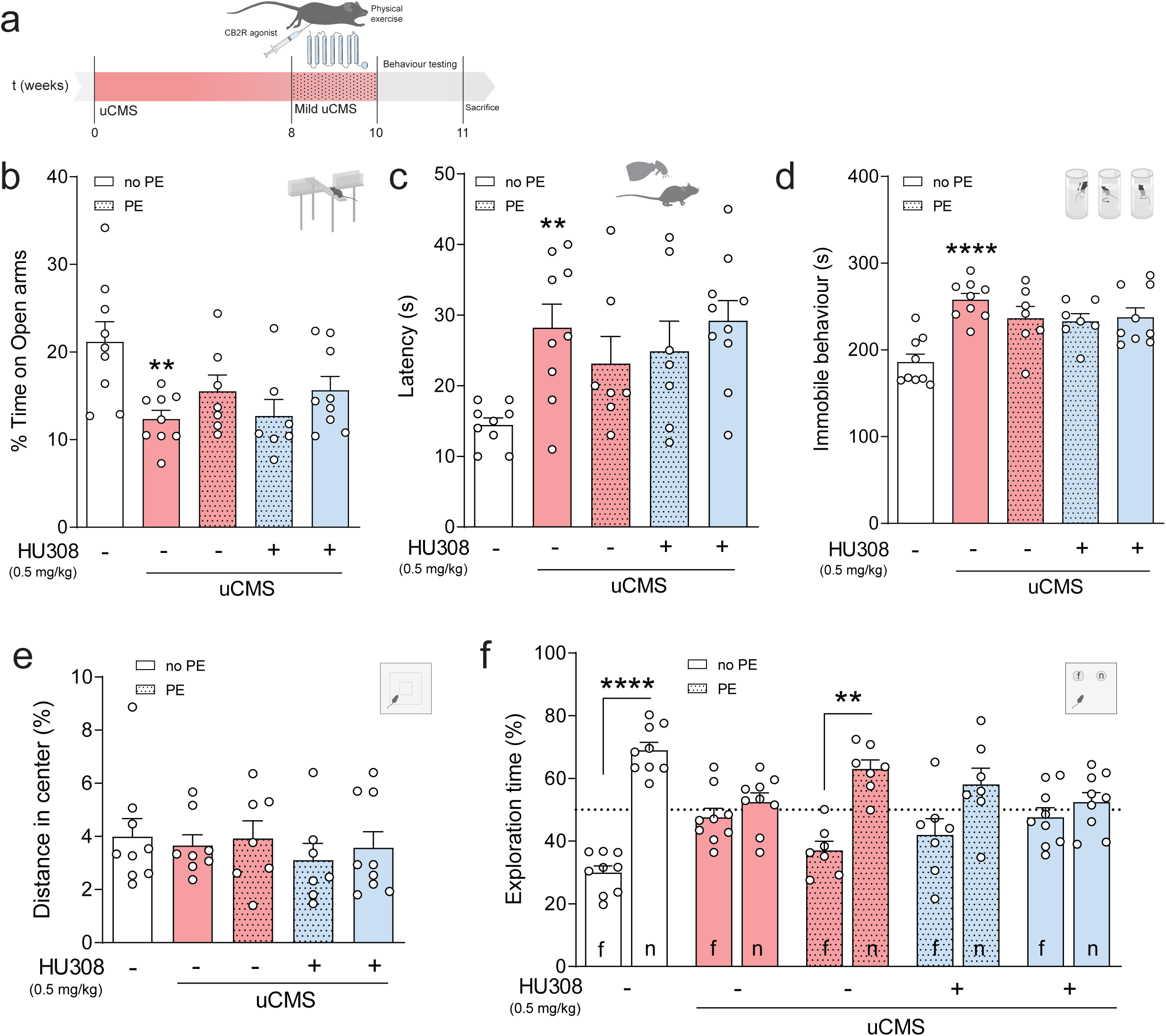

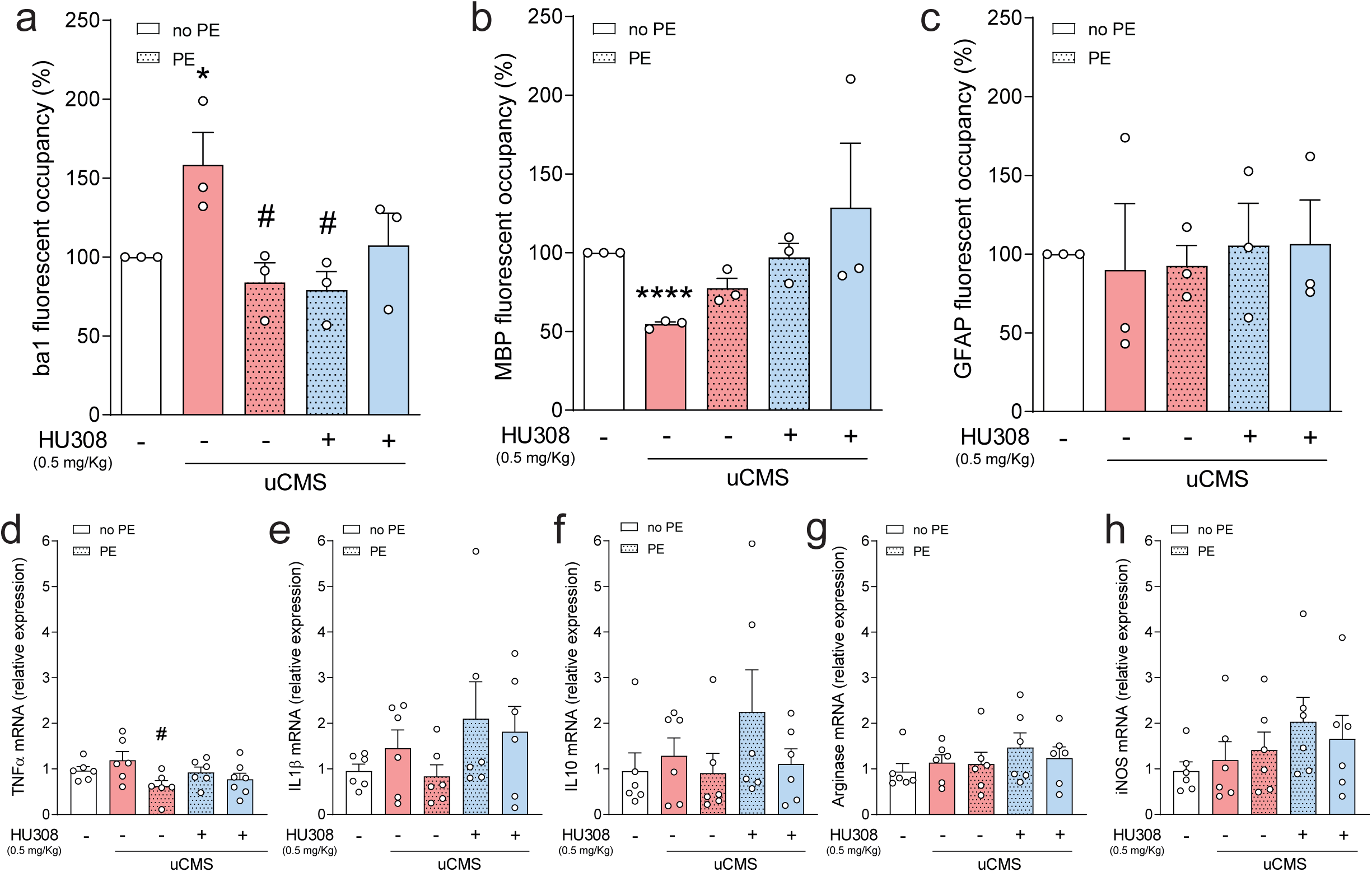

## Notes

### Competing Interest Statement

The authors have declared no competing interest.

## References

1. James SL, Abate D, Abate KH, Abay SM, Abbafati C, Abbasi N, et al. Global, regional, and national incidence, prevalence, and years lived with disability for 354 Diseases and Injuries for 195 countries and territories, 1990-2017: A systematic analysis for the Global Burden of Disease Study 2017. The Lancet. 2018;392:1789–1858.

2. Kraus C, Kadriu B, Lanzenberger R, Zarate CA, Kasper S. Prognosis and improved outcomes in major depression: a review. Translational Psychiatry 2019 9:1. 2019;9:1–17.

3. Rodrigues RS, Paulo SL, Moreira JB, Tanqueiro SR, Sebastião AM, Diógenes MJ, et al. Adult Neural Stem Cells as Promising Targets in Psychiatric Disorders. Stem Cells Dev. 2020;29:1099– 1117.

4. Toda T, Parylak SL, Linker SB, Gage FH. The role of adult hippocampal neurogenesis in brain health and disease. Mol Psychiatry. 2019;24:67–87.

5. Leschik J, Lutz B, Gentile A. Stress-Related Dysfunction of Adult Hippocampal Neurogenesis—An Attempt for Understanding Resilience? International Journal of Molecular Sciences 2021, Vol 22, Page 7339. 2021;22:7339.

6. Tartt AN, Mariani MB, Hen R, Mann JJ, Boldrini M. Dysregulation of adult hippocampal neuroplasticity in major depression: pathogenesis and therapeutic implications. Molecular Psychiatry 2022. 2022:1–11.

7. Boldrini M, Santiago AN, Hen R, Dwork AJ, Rosoklija GB, Tamir H, et al. Hippocampal granule neuron number and dentate gyrus volume in antidepressant-treated and untreated major depression. Neuropsychopharmacology. 2013;38.

8. Boldrini M, Galfalvy H, Dwork AJ, Rosoklija GB, Trencevska-Ivanovska I, Pavlovski G, et al. Resilience Is Associated With Larger Dentate Gyrus, While Suicide Decedents With Major Depressive Disorder Have Fewer Granule Neurons. Biol Psychiatry. 2019;85:850–862.

9. Snyder JS, Soumier A, Brewer M, Pickel J, Cameron HA. Adult hippocampal neurogenesis buffers stress responses and depressive behaviour. Nature. 2011;476:458–461.

10. Sahay A, Hen R. Adult hippocampal neurogenesis in depression. Nat Neurosci. 2007;10:1110– 1115.

11. Culig L, Surget A, Bourdey M, Khemissi W, le Guisquet AM, Vogel E, et al. Increasing adult hippocampal neurogenesis in mice after exposure to unpredictable chronic mild stress may counteract some of the effects of stress. Neuropharmacology. 2017;126:179–189.

12. Surget A, Tanti A, Leonardo ED, Laugeray A, Rainer Q, Touma C, et al. Antidepressants recruit new neurons to improve stress response regulation. Mol Psychiatry. 2011;16:1177–1188.

13. Planchez B, Surget A, Belzung C. Adult hippocampal neurogenesis and antidepressants effects. Curr Opin Pharmacol. 2020;50:88–95.

14. Schmidt HD, Duman RS. The role of neurotrophic factors in adult hippocampal neurogenesis, antidepressant treatments and animal models of depressive-like behavior. Behavioural Pharmacology. 2007;18:391–418.

15. Fogaça MV, Galve-Roperh I, Guimarães FS, Campos AC. Cannabinoids, neurogenesis and antidepressant drugs: is there a link? Curr Neuropharmacol. 2013;11:263–275.

16. Rodrigues RS, Lourenço DM, Paulo SL, Mateus JM, Ferreira MF, Mouro FM, et al. Cannabinoid Actions on Neural Stem Cells: Implications for Pathophysiology. Molecules. 2019;24:1350.

17. Vilar M, Mira H. Regulation of neurogenesis by neurotrophins during adulthood: Expected and unexpected roles. Front Neurosci. 2016;10.

18. deRoon-Cassini TA, Stollenwerk TM, Beatka M, Hillard CJ. Meet Your Stress Management Professionals: The Endocannabinoids. Trends Mol Med. 2020;26:953–968.

19. Gallego-Landin I, García-Baos A, Castro-Zavala A, Valverde O. Reviewing the Role of the Endocannabinoid System in the Pathophysiology of Depression. Front Pharmacol. 2021;12.

20. Morena M, Patel S, Bains JS, Hill MN. Neurobiological interactions between stress and the endocannabinoid system. Neuropsychopharmacology. 2016;41:80–102.

21. Jiang W, Zhang Y, Xiao L, van Cleemput J, Ji S-P, Bai G, et al. Cannabinoids promote embryonic and adult hippocampus neurogenesis and produce anxiolytic- and antidepressant-like effects. J Clin Invest. 2005;115:3104–3116.

22. Ishiguro H, Horiuchi Y, Tabata K, Liu QR, Arinami T, Onaivi ES. Cannabinoid CB2 receptor gene and environmental interaction in the development of psychiatric disorders. Molecules. 2018;23:1– 15.

23. Stempel AV, Stumpf A, Zhang HY, Özdoğan T, Pannasch U, Theis AK, et al. Cannabinoid Type 2 Receptors Mediate a Cell Type-Specific Plasticity in the Hippocampus. Neuron. 2016;90:795–809.

24. Bahi A, al Mansouri S, al Memari E, al Ameri M, Nurulain SM, Ojha S. β-Caryophyllene, a CB2 receptor agonist produces multiple behavioral changes relevant to anxiety and depression in mice. Physiol Behav. 2014;135:119–124.

25. Zoppi S, Madrigal JL, Caso JR, Garcia-Gutierrez MS, Manzanares J, Leza JC, et al. Regulatory role of the cannabinoid CB2 receptor in stress-induced neuroinflammation in mice. Br J Pharmacol. 2014;171:2814–2826.

26. García-Gutiérrez MS, Pérez-Ortiz JM, Gutiérrez-Adán A, Manzanares J. Depression-resistant endophenotype in mice overexpressing cannabinoid CB2 receptors. Br J Pharmacol. 2010;160:1773–1784.

27. García-Gutiérrez MS, Manzanares J. Overexpression of CB2 cannabinoid receptors decreased vulnerability to anxiety and impaired anxiolytic action of alprazolam in mice. J Psychopharmacol. 2011;25:111–120.

28. García-Gutiérrez MS, García-Bueno B, Zoppi S, Leza JC, Manzanares J. Chronic blockade of cannabinoid CB2 receptors induces anxiolytic-like actions associated with alterations in GABA(A) receptors. Br J Pharmacol. 2012;165:951–964.

29. Onaivi ES, Ishiguro H, Gong JP, Patel S, Meozzi PA, Myers L, et al. Brain neuronal CB2 cannabinoid receptors in drug abuse and depression: From mice to human subjects. PLoS One. 2008;3:e1640.

30. Hillard CJ, Liu Q. Endocannabinoid signaling in the etiology and treatment of major depressive illness. Curr Pharm Des. 2014;20:3795–3811.

31. Saraulli D, Costanzi M, Mastrorilli V, Farioli-Vecchioli S. The Long Run: Neuroprotective Effects of Physical Exercise on Adult Neurogenesis from Youth to Old Age. Curr Neuropharmacol. 2017;15:519.

32. Cotman CW, Berchtold NC, Christie LA. Exercise builds brain health: key roles of growth factor cascades and inflammation. Trends Neurosci. 2007;30:464–472.

33. Castrén E, Võikar V, Rantamäki T. Role of neurotrophic factors in depression. Curr Opin Pharmacol. 2007;7:18–21.

34. van Praag H. Neurogenesis and Exercise: Past and Future Directions. Neuromolecular Med. 2008;10:128–140.

35. Wang L, Chang X, She L, Xu D, Huang W, Poo M -m. Autocrine Action of BDNF on Dendrite Development of Adult-Born Hippocampal Neurons. Journal of Neuroscience. 2015;35:8384–8393.

36. Choi SH, Li Y, Parada LF, Sisodia SS. Regulation of hippocampal progenitor cell survival, proliferation and dendritic development by BDNF. Mol Neurodegener. 2009;4:52.

37. Troubat R, Barone P, Leman S, Desmidt T, Cressant A, Atanasova B, et al. Neuroinflammation and depression: A review. European Journal of Neuroscience. 2021;53:151–171.

38. Forteza F, Giorgini G, Raymond F. Neurobiological Processes Induced by Aerobic Exercise through the Endocannabinoidome. Cells. 2021;10.

39. Charytoniuk T, Zywno H, Konstantynowicz-Nowicka K, Berk K, Bzdega W, Chabowski A. Can physical activity support the endocannabinoid system in the preventive and therapeutic approach to neurological disorders? Int J Mol Sci. 2020;21:1–16.

40. Raichlen D a, Foster AD, Seillier A, Giuffrida A, Gerdeman GL. Exercise-induced endocannabinoid signaling is modulated by intensity. Eur J Appl Physiol. 2013;113:869–875.

41. Wang H, Han J. The endocannabinoid system regulates the moderate exercise-induced enhancement of learning and memory in mice. Journal of Sports Medicine and Physical Fitness. 2020;60:320–328.

42. Meyer JD, Crombie KM, Cook DB, Hillard CJ, Koltyn KF. Serum Endocannabinoid and Mood Changes after Exercise in Major Depressive Disorder. Med Sci Sports Exerc. 2019;51:1909–1917.

43. Marin Bosch B, Bringard A, Logrieco MG, Lauer E, Imobersteg N, Thomas A, et al. A single session of moderate intensity exercise influences memory, endocannabinoids and brain derived neurotrophic factor levels in men. Scientific Reports 2021 11:1. 2021;11:1–14.

44. Ferreira FF, Ribeiro FF, Rodrigues RS, Sebastião AM, Xapelli S. Brain-derived neurotrophic factor (BDNF) role in cannabinoid-mediated neurogenesis. Front Cell Neurosci. 2018;12:1–16.

45. Ribeiro MA, Aguiar RP, Scarante FF, Fusse EJ, W de Oliveira RM, Guimaraes FS, et al. Spontaneous Activity of CB2 Receptors Attenuates Stress-Induced Behavioral and Neuroplastic Deficits in Male Mice. Front Pharmacol. 2022;12.

46. Willner P. The chronic mild stress (CMS) model of depression: History, evaluation and usage. Neurobiol Stress. 2017;6:78–93.

47. Surget A, Saxe M, Leman S, Ibarguen-Vargas Y, Chalon S, Griebel G, et al. Drug-Dependent Requirement of Hippocampal Neurogenesis in a Model of Depression and of Antidepressant Reversal. Biol Psychiatry. 2008;64:293–301.

48. Monteiro S, Roque S, de Sá-Calçada D, Sousa N, Correia-Neves M, Cerqueira JJ, et al. An efficient chronic unpredictable stress protocol to induce stress-related responses in C57BL/6 mice. Front Psychiatry. 2015;6.

49. Patki G, Li L, Allam F, Solanki N, Dao AT, Alkadhi K, et al. Moderate treadmill exercise rescues anxiety and depression-like behavior as well as memory impairment in a rat model of posttraumatic stress disorder. Physiol Behav. 2014;130:47–53.

50. Lad HV, Liu L, Paya-Cano JL, Parsons MJ, Kember R, Fernandes C, et al. Behavioural battery testing: evaluation and behavioural outcomes in 8 inbred mouse strains. Physiol Behav. 2010;99:301–316.

51. Mouro FM, Batalha VL, Ferreira DG, Coelho JE, Baqi Y, Müller CE, et al. Chronic and acute adenosine A2Areceptor blockade prevents long-term episodic memory disruption caused by acute cannabinoid CB1receptor activation. Neuropharmacology. 2017;117:316–327.

52. Porsolt RD, Anton G, Blavet N, Jalfre M. Behavioural despair in rats: A new model sensitive to antidepressant treatments. Eur J Pharmacol. 1977;47:379–391.

53. Walf AA, Frye CA. The use of the elevated plus maze as an assay of anxiety-related behavior in rodents. Nat Protoc. 2007;2:322–328.

54. Hoffman KL. What can animal models tell us about depressive disorders? Modeling Neuropsychiatric Disorders in Laboratory Animals, Woodhead Publishing; 2016. p. 35–86.

55. Ferreiro E, Lanzillo M, Canhoto D, Carvalho da Silva AM, Mota SI, Dias IS, et al. Chronic hyperglycemia impairs hippocampal neurogenesis and memory in an Alzheimer’s disease mouse model. Neurobiol Aging. 2020;92:98–113.

56. Alves ND, Patrício P, Correia JS, Mateus-Pinheiro A, Machado-Santos AR, Loureiro-Campos E, et al. Chronic stress targets adult neurogenesis preferentially in the suprapyramidal blade of the rat dorsal dentate gyrus. Brain Struct Funct. 2018;223:415–428.

57. Tanti A, Belzung C. Neurogenesis along the septo-temporal axis of the hippocampus: are depression and the action of antidepressants region-specific? Neuroscience. 2013;252:234–252.

58. Hawley DF, Leasure JL. Region-specific response of the hippocampus to chronic unpredictable stress. Hippocampus. 2012;22:1338–1349.

59. Ribeiro FF, Ferreira F, Rodrigues RS, Soares R, Pedro DM, Duarte-Samartinho M, et al. Regulation of hippocampal postnatal and adult neurogenesis by adenosine A2A receptor: Interaction with brain-derived neurotrophic factor. Stem Cells. 2021;39:1362–1381.

60. van Praag H, Christie BR, Sejnowski TJ, Gage FH. Running enhances neurogenesis, learning, and long-term potentiation in mice. Proc Natl Acad Sci U S A. 1999;96:13427–13431.

61. Gubert C, Hannan AJ. Exercise mimetics: harnessing the therapeutic effects of physical activity. Nat Rev Drug Discov. 2021;20:862–879.

62. Sohroforouzani AM, Shakerian S, Ghanbarzadeh M, Alaei H. Effect of forced treadmill exercise on stimulation of BDNF expression, depression symptoms, tactile memory and working memory in LPS-treated rats. Behavioural Brain Research. 2022;418.

63. Hillard CJ. Circulating Endocannabinoids: From Whence Do They Come and Where are They Going? Neuropsychopharmacology. 2018;43:155–172.

64. Hill MN, Patel S, Campolongo P, Tasker JG, Wotjak CT, Bains JS. Functional interactions between stress and the endocannabinoid system: from synaptic signaling to behavioral output. J Neurosci. 2010;30:14980–14986.

65. Heyman E, Gamelin FX, Goekint M, Piscitelli F, Roelands B, Leclair E, et al. Intense exercise increases circulating endocannabinoid and BDNF levels in humans-Possible implications for reward and depression. Psychoneuroendocrinology. 2012;37:844–851.

66. Hill MN, Titterness AK, Morrish AC, Carrier EJ, Lee TT-Y, Gil-mohapel J, et al. Endogenous cannabinoid signaling is required for voluntary exercise-induced enhancement of progenitor cell proliferation in the hippocampus. Hippocampus. 2010;523:513–523.

67. Burstein O, Shoshan N, Doron R, Akirav I. Cannabinoids prevent depressive-like symptoms and alterations in BDNF expression in a rat model of PTSD. Prog Neuropsychopharmacol Biol Psychiatry. 2018;84:129–139.

68. Li Y, Kim J. CB2 Cannabinoid Receptor Knockout in Mice Impairs Contextual Long-Term Memory and Enhances Spatial Working Memory. Neural Plast. 2015;2016.

69. Aso E, Juvés S, Maldonado R, Ferrer I. CB2 Cannabinoid Receptor Agonist Ameliorates Alzheimer-Like Phenotype in AβPP/PS1 Mice. Journal of Alzheimer’s Disease. 2013;35:847–858.

70. Nollet M, Gaillard P, Tanti A, Girault V, Belzung C, Leman S. Neurogenesis-independent antidepressant-like effects on behavior and stress axis response of a dual orexin receptor antagonist in a rodent model of depression. Neuropsychopharmacology. 2012;37:2210–2221.

71. Jayatissa MN, Bisgaard C, Tingström A, Papp M, Wiborg O. Hippocampal cytogenesis correlates to escitalopram-mediated recovery in a chronic mild stress rat model of depression. Neuropsychopharmacology. 2006;31:2395–2404.

72. Tanti A, Rainer Q, Minier F, Surget A, Belzung C. Differential environmental regulation of neurogenesis along the septo-temporal axis of the hippocampus. Neuropharmacology. 2012;63:374–384.

73. Nishijima T, Kawakami M, Kita I. Long-Term Exercise Is a Potent Trigger for ΔFosB Induction in the Hippocampus along the dorso–ventral Axis. PLoS One. 2013;8:e81245.

74. Castilla-Ortega E, Rosell-Valle C, Pedraza C, Rodríguez de Fonseca F, Estivill-Torrús G, Santín LJ. Voluntary exercise followed by chronic stress strikingly increases mature adult-born hippocampal neurons and prevents stress-induced deficits in ‘what-when-where’ memory. Neurobiol Learn Mem. 2014;109:62–73.

75. Snyder JS, Glover LR, Sanzone KM, Kamhi JF, Cameron H a. The effects of exercise and stress on the survival and maturation of adult-generated granule cells. Hippocampus. 2009;19:898–906.

76. Mensching L, Djogo N, Keller C, Rading S, Karsak M. Stable adult hippocampal neurogenesis in cannabinoid receptor CB2 deficient mice. Int J Mol Sci. 2019;20.

77. Berdugo-Vega G, Lee CC, Garthe A, Kempermann G, Calegari F. Adult-born neurons promote cognitive flexibility by improving memory precision and indexing. Hippocampus. 2021;31:1068– 1079.

78. Wu M v., Sahay A, Duman RS, Hen R. Functional differentiation of adult-born neurons along the septotemporal axis of the dentate gyrus. Cold Spring Harb Perspect Biol. 2015;7.

79. Tuncdemir SN, Lacefield CO, Hen R. Contributions of adult neurogenesis to dentate gyrus network activity and computations. Behavioural Brain Research. 2019;374.

80. Heine VM, Maslam S, Zareno J, Joëls M, Lucassen PJ. Suppressed proliferation and apoptotic changes in the rat dentate gyrus after acute and chronic stress are reversible. European Journal of Neuroscience. 2004;19:131–144.

81. Schoenfeld TJ, Rada P, Pieruzzini PR, Hsueh B, Gould E. Physical Exercise Prevents Stress-Induced Activation of Granule Neurons and Enhances Local Inhibitory Mechanisms in the Dentate Gyrus. Journal of Neuroscience. 2013;33:7770–7777.

82. Sun D, Milibari L, Pan JX, Ren X, Yao LL, Zhao Y, et al. Critical Roles of Embryonic Born Dorsal Dentate Granule Neurons for Activity-Dependent Increases in BDNF, Adult Hippocampal Neurogenesis, and Antianxiety-like Behaviors. Biol Psychiatry. 2021;89:600–614.

83. Mograbi KDM, Suchecki D, da Silva SG, Covolan L, Hamani C. Chronic unpredictable restraint stress increases hippocampal pro-inflammatory cytokines and decreases motivated behavior in rats. Stress. 2020;23:427–436.

84. Kreisel T, Frank MG, Licht T, Reshef R, Ben-Menachem-Zidon O, Baratta M v., et al. Dynamic microglial alterations underlie stress-induced depressive-like behavior and suppressed neurogenesis. Mol Psychiatry. 2014;19:699–709.

85. Mee-inta O, Zhao Z-W, Kuo Y-M. Physical Exercise Inhibits Inflammation and Microglial Activation. Cells 2019, Vol 8, Page 691. 2019;8:691.

86. dos Santos RS, Sorgi CA, Peti APF, Veras FP, Faccioli LH, Galdino G. Involvement of Spinal Cannabinoid CB 2 Receptors in Exercise-Induced Antinociception. Neuroscience. 2019;418:177– 188.

87. Komorowska-Müller JA, Schmöle AC. CB2 receptor in microglia: The guardian of self-control. Int J Mol Sci. 2021;22:1–27.

88. Valencia-Sanchez S, Nava-Castro KE, Palacios-Arreola MI, Prospero-Garcia O, Morales-Montor J, Drucker-Colin R. Chronic exercise modulates the cellular immunity and its cannabinoid receptors expression. PLoS One. 2019;14:e0220542.

89. Anacker C, Luna VM, Stevens GS, Millette A, Shores R, Jimenez JC, et al. Hippocampal neurogenesis confers stress resilience by inhibiting the ventral dentate gyrus. Nature. 2018;559:98– 102.

90. Bolognini D, Cascio MG, Parolaro D, Pertwee RG. AM630 behaves as a protean ligand at the human cannabinoid CB2 receptor. Br J Pharmacol. 2012;165:2561.

91. Kenakin T. Inverse, protean, and ligand-selective agonism: matters of receptor conformation. FASEB J. 2001;15:598–611.

