## Supplementary material for "Cannabinoid type 2 receptor inhibition enhances the antidepressant and proneurogenic effects of physical exercise after chronic stress"

**Supplementary Materials and Methods**

**Experimental design and interventional procedures**

Given the wide range of doses (0.01 – 20 mg/kg) used in studies addressing the effects of pharmacological CB2R modulators, CB2R dose-response experiments were performed. Briefly, subchronic administration of CB2R ligands (CB2R agonist (HU308, Tocris, Bristol, UK) or inverse agonist (AM630, Tocris, Bristol, UK) at 0.5 and 5 mg/kg) was performed for 2 weeks followed by behavioural assessment. For stress experiments, an unpredictable chronic mild stress (uCMS) protocol was validated in our laboratory in a pilot study and applied for an 8-week period (**Fig. S2a,b**). Briefly, this stress paradigm induces depressive-like behaviour, anxiety-like phenotype and cognitive deficits in animals through random and unpredictable exposure to a wide range of different mild stressors. Specifically, in our modified uCMS protocol, animals were exposed to three stressors per day (exact uCMS schedule in **Fig. S2b**), which included distinct daily and night stressors such as spatial confinement (individual or group confinement), cage shaking, exposure to stroboscopic lights, overnight placement in a tilted cage (45°) or damp bedding, reversed light/dark cycle for 48 h. Food or water deprivation was not performed. Following this 8-week uCMS protocol, a 2-week uCMS milder protocol was implemented in which both stressors (two stressors per day) and interventional procedures (CB2R pharmacological treatments and PE) were applied. During this period, mice were pharmacologically treated with either aforementioned CB2R ligands (at 0.5 mg/kg) or vehicle treatment (saline solution, 0.9% NaCl) intraperitoneally (i.p.). In some groups, a 2-week forced physical exercise (PE) protocol was performed consisting of 30 min/daily sessions at a 10 m/min speed in the first week and at a 15m/min speed in the second week. Experimental groups submitted to PE ran on average 300 and 450 meters/day in the first and second weeks of treatment, respectively, and no differences were observed between groups (data not shown). For all experiments, husbandry and physical health measures were routinely monitored. Body weight, evaluated weekly, is reported as a percentage of weight change relative to baseline measures (taken on the first week of handling) (**Fig. S2c**). Four weeks (28 days) before sacrifice, all animals received i.p. injections of BrdU (50 mg/kg; Sigma Aldrich, MO, USA) during 5 consecutive days to label newly generated adult-born neurons.

**Behavioural assessment**

Behavioural tests were performed during one week after interventional procedures (one test per day, with the exception of EPM and NOR habituation day 3 that were performed in the same day) (**Fig. S2a,b**). Animals were allowed to habituate to the experimental room 1h prior to testing. The order in which each assay was carried out was based on previous work, starting with the tasks requiring maximal levels of novelty (i.e., OF, EPM, NOR) and ending with the most stressful ones (i.e., FST, SST). Between tests and trials, the corresponding apparatus and/or objects were carefully cleaned with a 30% ethanol solution to erase any olfactory cues. All assays were videorecorded for posterior manual or software-driven analyses. OF and EPM data was videorecorded and analysed using the SMART® v2.5 video-tracking software (Panlab, Harvard Apparatus, Barcelona, Spain). NOR, FST and SST assays were videorecorded and subsequently manually analysed by a trained experimenter blind to experimental conditions using Solomon Coder beta 17.03.22 (András Péter, Milan, Italy) behaviour coding software.

**Open Field (OF) test**

To assess locomotor, exploratory and anxiety-like behaviour, the OF test was performed. Briefly, each animal was placed individually in the centre of an empty square box (60x60x40 cm), virtually divided in three squares at a similar distance from each other, and allowed to freely explore it for 5 min, without prior habituation. Total distance travelled and average velocity of the animals were considered measures of locomotor capacity, and the percentage of distance travelled in the centre of the arena was considered as measurement of anxiety-like behaviour. This test was also used as part of the first day of the habituation phase for the NOR.

**Elevated-plus Maze (EPM)**

To assess anxiety-like behaviour, the EPM test was also performed. Briefly, animals were individually placed in the centre of an EPM apparatus, with two opposite open arms (50x10 cm) and two closed arms (50x10x30 cm) raised 50 cm above the floor and dimly illuminated, facing an open arm, and left to explore the maze for 5 minutes, with the number of entries and time spent in the open arms being recorded.

**Novel Object Recognition (NOR)**

To evaluate long-term memory performance, NOR was performed. It was composed of a habituation phase (3 consecutive days) in which animals freely explored the empty OF arena for 15 min, followed by a familiarization phase (in the fourth day) in which animals were presented with two to-be-familiarized objects (familiar objects) for 5 min. After 24h, animals returned to the arena for the test phase, where it was presented with one of the previously explored objects and a novel object, for 5 min. The objects consisted of different wooden dolls, proportional to the size of mice, randomized and used interchangeably as familiar and novel objects between trials. Exploration was scored whenever the animal touched an object with its forepaws and/or bit, sniffed or reared towards the object. Running around the object, standing next to it with its head facing away or climbing it was excluded. The total exploration time of both objects in the familiarization or test phase was used as a measure of exploratory drive. Exploration time for each object in the training phase was used to exclude preference for one of the familiar objects, whereas in the test phase it was used as a measure of memory performance (quantified as the percentage time spent exploring each object over the total time spent exploring both objects).

**Forced-swimming test (FST)**

To assess adaptive behaviour, i.e., the ability to cope with an inescapable stressor, the forced swimming test or Porsolt swim test was performed as a measure of depressive-like behaviour. Briefly, each animal was individually placed in a glass cylinder filled with water (23 °C; depth 30 cm) for 6 min. All sessions were videorecorded, and time spent engaging in three distinct behaviours (climbing, swimming, immobility) during the last 5 min of the test was quantified. Immobility time was considered as a measure of coping behaviour. Mice were considered immobile when all active behaviours (struggling, swimming, and jumping) were ceased. For immobility, the animals had to remain passively floating or making minimal movements required to maintain the nostrils above water. After each trial, animals were dried and placed under a heating lamp for at least 10 minutes, and the cylinder water was changed.

**Sucrose splash test (SST)**

To evaluate self-caring and motivational behaviour, SST was performed. Briefly, each animal was sprayed with a 10 % sucrose solution on the dorsal coat in a separate test cage under red light. Grooming behaviours were stimulated by the palatability of the solution. After spraying the animals, behaviour was videorecorded for 5 min and latency time, grooming frequency and duration, considered measures of anhedonic-like behaviour, were quantified.

**Animal sacrifice and tissue processing**

Animals were anesthetized with sodium pentobarbital (70 mg/kg i.p.) and transcardially perfused for 10 min with phosphate-buffered saline (PBS) (137 mM NaCl, 2.7 mM KCl, 1.8 mM KH_2_PO_4_ and 10 mM Na_2_HPO_4_.2H_2_O, pH = 7.4), except for electrophysiological experiments (see below). Brains were carefully removed from the skull and left hemispheres were postfixed in 4 % paraformaldehyde (PFA) for 24 h at 4 ºC followed by cryoprotection in sequential 15 % and 30% sucrose in PBS (4 °C). Right hemispheres were quickly sectioned at 450 μm (McIlwainTM Tissue Chopper, Campden Instruments, Loughborough, UK) and regions of interest (e.g., DG) were quickly microdissected and subsequently flash-frozen in liquid nitrogen and stored at -80 ºC. Left hemispheres were then gelatine-embedded and coronally sectioned at 40 μm (cryostat Leica CM3050 S, Leica Biosystems, Germany). Hippocampi-containing sections were collected, in ten series, each one comprising an anterior-posterior reconstruction of the hippocampus (sections separated by 400 μm) and stored at -20 °C in anti-freezing medium (30 % glycerol, 30 % ethylene glycol, phosphate buffer 0.1 M (8.9 % Na_2_HPO_4_.2H_2_O, 7.8 % NaH_2_PO_4_.2H_2_O, pH = 7.4)).

**Tissue processing and immunostainings**

Degelatinization was achieved by sequentially immersing free-floating brain slices in PBS at 37 °C for 8 minutes. In all immunostainings related to AHN assessment (except for BrdU/DCX immunostainings), an antigen retrieval step was primarily performed using citrate buffer 1X (pH = 6) at 80ºC for 25 min followed by 25 min at room temperature (RT). For BrdU/DCX or BrdU/NeuN immunostainings, slices were incubated during 20 min in 2 N HCl at 37 °C in order to denature DNA to expose BrdU epitopes for antibody binding, followed by acid pH neutralization with borate buffer 1 M (pH 8.5) for (2 x 5 min), and several washes with PBS (3 x 8 min). Brain slices were incubated for 1h at RT with blocking and permeabilization solution of PBS containing 6 % bovine serum albumin (BSA) and 1 % Triton X-100. In the same blocking solution, brain slices were double stained for BrdU (#M0744; mouse, 1:500; Agilent, CA, USA) and DCX (#sc-8066; goat, 1:500; Santa Cruz Biotechnologies, CA, USA) or BrdU (#ab6326; rat, 1:500; Abcam, UK) and NeuN (#12943S; rabbit, 1:500; Cell Signalling Technology, MO, USA) for two overnights in a humidified chamber (4°C). For Ki67/DCX immunostainings, brain slices were incubated for 1h at RT with blocking and permeabilization solution of PBS containing 3% BSA and 0.2% Triton X-100 right after the antigen retrieval step. In the same blocking solution, brain slices were double stained for Ki67 (#ab16667; rabbit, 1:500; Abcam, UK) and DCX (#326006; chicken, 1:750; Synaptic Systems, Germany) for two overnights in a humidified chamber (4 °C). In all immunostainings, washing with PBS (3 x 8 min) was performed following incubation with primary antibodies, and incubation in blocking solution containing the appropriate secondary antibodies (Alexa Fluor 488 goat anti-mouse polyclonal, #A-11001; 1:500; Alexa Fluor 488 goat anti-chicken, #A-11039; 1:1000; Alexa Fluor 488 goat anti-rat, #A-11006; 1:1000; Alexa Fluor 568 donkey anti-goat, #A-11057; 1:500 and Alexa Fluor 568 goat anti-rabbit, #A-11011; 1:1000; all from Thermo Fisher Scientific, MA, USA) and DAPI (1μg/ml, Sigma-Aldrich, MO, USA) for 2h at RT followed. After a final wash with PBS (3 x 8 min), slices were mounted onto microscope slides (Superfrost™ Plus, Thermo Fisher Scientific, MA, USA) with Mowiol fluorescent medium and glass coverslips.

For other immunostainings (Iba1/GFAP/MBP), antigen retrieval was not performed. Brain slices were permeabilized with 0.2 % Triton X-100 in PBS for 10 min and further incubated with a blocking solution of 5 % BSA, 5 % fetal bovine serum (FBS) and 0.1 % Triton X-100 for 1 h at RT. In the same blocking solution, brain slices were triple stained for Iba1 (#010-19741; rabbit, 1:250; FUJIFILM Wako Chemicals Europe GmbH, Germany), GFAP (#NCL-GFAP-GA5; mouse, 1:100; Novocastra, Leica Biosystems, Germany) and MBP (#MCA409S; rat, 1:200; Bio-Rad Laboratories, CA, USA) for two overnights in a humidified chamber (4°C). Washing with PBS (3 x 10 min) was followed by incubation in blocking solution containing the appropriate secondary antibodies (Alexa Fluor 488 goat anti-rabbit, #A-11008; 1:500; Thermo Fisher Scientific, MA, USA; and Alexa Fluor 568 goat anti-rat, #A-11077; 1:500; Thermo Fisher Scientific, MA, USA; and Alexa Fluor 647 goat anti-mouse, #A-21235; 1:500) and DAPI (1μg/ml, Sigma-Aldrich, MO, USA) for 2h at RT followed. Afterwards, washing with PBS (3 x 10 min) was followed by incubation with DAPI (1:1000) for 5 min. Finally, the slices were washed three times for 5 minutes each with PBS and then, mounted onto microscope slides using Fluoromount-G (Southern Biotech, AL, USA).

**Image acquisition and stereology**

For stereological cell counting, images of the immunostained coronal sections, containing dorsal and ventral hippocampi, were captured using a Zeiss Cell Observer SD spinning disk confocal microscope (Carl Zeiss, Oberkochen, Germany) with a 40x water immersion objective (40x/1.1, NA 1.1). All images were acquired throughout the entire thickness of the corresponding coronal section (Z-stack). The quantification of each cell population (Ki67+, DCX+, Ki67+DCX+ or BrdU+NeuN+ cells) was performed in hippocampal slices of one-in-ten series, representing the reconstruction of the whole hippocampus throughout the rostro-caudal extent of the brain (from -0.94 to -4.04 mm of bregma), with slices being apart from 400 μm. At least eight DG sections per animal (4 sections for dorsal hippocampus [-1.06 to -2.06 mm of bregma] and 4 sections for ventral hippocampus [-3.08 to -4.04 mm of bregma]) were included in the analysis. The density of each cell population in the DG (only SGZ and GCL, not hilus, were considered) was achieved by manual counting immunolabelled positive cells across Z-stack, in an unbiased and blinded stereological fashion, and defined by the ratio of the total number of cells and the corresponding volume of DG sections (total, dorsal (dHip) or ventral (vHip)).

Areas and volumes of each DG section were obtained from serial sections with 400 μm rostro-caudal intervals along the dorsal and ventral hippocampi (8-10 sections per animal), using immunofluorescent staining for DAPI and Zeiss ZEN 2.1 software (blue edition). The area of each section was measured at maximum intensity projection of each Z-stack, using DAPI staining, and the volume of DG sections for each animal was calculated by multiplying the sum of the areas by the thickness of each section (40 μm). The total DG volume (mm^3^) for each animal as well as dorsal or ventral volumes, were extrapolated by multiplying the sum of the areas, or corresponding areas, by the distance between slices (400 μm) and no volumetric differences were found between tested conditions. Cell densities are reported as number of cells per total estimated DG volume or per dorsal or ventral estimated volumes (mm^3^) (**Table S2**).

The impact of different interventional procedures (e.g., uCMS, PE, CB2R ligands) on the subregional distribution of each cell population of dorsal or ventral hippocampi was calculated by determining the percentage of cells in each subregion. To assess the contribution of specific DG subregions (in corresponding dorsal and ventral DG) to observed effects, the subregional percentage of immunopositive cells was calculated and compared between the different DG subregions, including supra vs infrapyramidal blades and SGZ vs GCL. Relative proportions were determined by taking into consideration the percentages of immunopositive cells in each region in function of the corresponding area as follows: percentage of immunopositive cells in subregion 1 / (percentage of immunopositive cells in the subregion 1 + percentage of immunopositive cells in subregion 2). For example, the regional distribution of BrdU+NeuN+ cells in the SGZ subregion of the dorsal DG suprapyramidal blade (in comparison to the GCL subregion of the same area) was determined as follows: percentage of supraSGZ BrdU+NeuN+ cells / (percentage supraSGZ BrdU+NeuN+ cells + supraGCL BrdU+NeuN+ cells). Moreover, DCX+ cells were classified based on their level of maturation (assessed by morphological features and localization) and divided into two subpopulations: the neuronal progenitors (“DCX+ neuronal progenitors”), which are type 2b and type 3 cells, located in the SGZ or immediately adjacent, displaying no processes or short processes parallel to the GCL; and the immature neurons (“DCX+ immature neurons”) located immediately outside the SGZ or in the middle third part of the GCL, displaying a branched dendritic extension radially crossing the GCL.

For experiments assessing the expression levels of microglia (Iba1), astrocytes (pan GFAP) and oligodendrocytes (MBP), fluorescent images were obtained by confocal microscopy using Zeiss Cell Observer SD spinning disk confocal microscope with Zeiss ZEN 2.1 software (black edition). For each independent experiment at least 3 hippocampal slices (at least 1 dorsal and 1 ventral) were used. The range of the slice was determined using Z-stack imaging at 1-µm intervals, and a series of images derived from was merged Z-stack imaging (maximum intensity projections) were analysed. The percentage of fluorescent occupancy for each antibody was measured in the DG granular layer (results used for **Figs. 4**, **5**, and **S4**). A cut off intensity threshold, i.e., a minimum intensity due to specific staining above background values, was defined for each region of interest (ROI) and maintained across different experiments. The percentage of fluorescent occupancy of Iba1, GFAP and MBP above set threshold was measured in each ROI automatically using Fiji software and normalized to control (CTRL) levels (set to 100%).

**Biochemical and molecular analyses**

Serum corticosterone levels were measured in all groups (n = 8-10 per group) using a commercially available enzyme-linked immunosorbent assay (ELISA) kit (Abcam, UK), according to the manufacturer’s instructions. Briefly, sampling (submandibular venepuncture) was performed between 8-9 am at the end of the stress protocol and obtained sera was stored at -20 ºC. Each sample was diluted 1:100 in ELISA buffer. Next, two corticosterone standards were made from a 100 ng/ml corticosterone standard solution using a serial dilution method. Blank samples, standards and diluted samples were then transferred directly onto the ELISA plate pre-coated with corticosterone antibody. Biotinylated corticosterone protein was subsequently added, and the plate incubated for 2 h at RT on a plate shaker moving at 220 rpm. After washing, conjugate was added for 30 min followed by a chromogen substrate solution and the plate was incubated at RT for 20 min, before being terminated with a stop solution. Absorbance was immediately read at a wavelength of 450 nm using multimode microplate reader (Infinite M200, Tecan) and readings at 570 nm were used to correct for optical imperfections. Results are expressed as ng (of corticosterone) per ml (of serum).

Total RNA was isolated using RiboZol™ reagent method, according to the manufacture’s guidelines (VWR Life Science, USA). RNA concentration and purity was checked using NanoDrop ND-100 Spectrophotometer (NanoDrop Techonologies, Wilmington, DE, USA). A total of 400 ng RNA was reversibly transcribed into cDNA using Xpert cDNA Synthesis Mastermix kit (GRiSP) and cDNA was amplified by semi-quantitative RT-PCR (qRT-PCR) on a 7300 Real-Time PCR System (Applied Biosystem, Madrid, Spain) by the excitation and emission of Xpert Fast SYBR Mastermix (GRiSP). The cycle conditions were previously optimized: 50 ºC for 2 min, 95 ºC for 10 min followed by 40 cycles at 95 ºC for 15 seconds and 62 ºC for 1 min. The PCR was performed in 384-well plates, with duplicates for each sample. The sequences used as primers are listed in **Table S3**. All primers were purchased from Thermo Fisher Scientiﬁc, MA, USA. Relative mRNA concentration was measured using the ∆∆Ct comparative method, where ∆Ct was calculated by the difference between the Ct value of the gene of interest and the mean Ct value of β-actin. Then, ∆∆Ct value of one sample was achieved by the difference between its ∆Ct value and the ∆Ct value of the sample chosen as reference, in our case the CTRL group. Results are presented as relative expression to the standard gene.

Western blotting analysis was used to assess CB2R protein levels from extracted DG tissue. Samples were collected into a urea 8 M (in Tris-HCl 1M, pH = 8) and 1 % SDS (in H_2_O) solution (1:1 ratio), containing a protease inhibitor cocktail tablet (Roche, Switzerland) (pH 7.4 at 4°C), and sonicated for 2 min. Samples were centrifuged for 10 min at 13,000 g and 4 °C. The protein concentration of the supernatant was determined using the BCA method. Samples with equal amounts of protein (50 μg) samples were treated with SDS-PAGE sample buffer (5x sample buffer: 350 mM Tris, 30 % glycerol, 10 % SDS, 600 mM dithiothreitol, and 0.012 % bromophenol blue, pH = 6.8). Protein running was performed in SDS-PAGE gels (10 % acrylamide/bisacrylamide for resolving and 5 % for stacking gels), transferred to polyvinylidene difluoride (PVDF) membranes, and blocked for 1h at room temperature with 5 % milk in TBS-T (Tris buffer saline with 0.1 % Tween-20, 200 nM Tris, 1.5 M NaCl). Incubation with the primary antibody anti-CB2R (#sc-293188; mouse, 1:1000; Santa Cruz Biotechnologies, CA, USA) was performed overnight at 4 °C followed by incubation with the secondary antibody (#172-1011; goat anti-mouse IgG (H+L)-HRP conjugate, 1:5000; Bio-Rad Laboratories, CA, USA) for 1 h at room temperature. For the loading control clocking for 1h at room temperature with 3% BSA in TBS-T was used followed by primary antibody against GAPDH (#AM4300; mouse, 1:5000; Thermo Fisher Scientific, MA, USA) and secondary antibody (#172-1011; goat anti-mouse IgG (H+L)-HRP conjugate, 1:5000; Bio-Rad Laboratories, CA, USA) incubations. Membranes were then incubated with ECL plus western blotting detection reagent (GE Healthcare, Buckinghamshire, UK) and protein immunoreactive bands were visualized with ChemiDoc Imaging System (model XRS+, Bio-Rad Laboratories, CA, USA). Optical density of protein bands was determined with Fiji software. Results were normalized to the expression of GAPDH.

**Electrophysiological recordings**

Animals were anesthetized with sodium pentobarbital (70 mg/kg i.p.) and transcardially perfused for 3 min with oxygenated (95% O_2_, 5% CO_2_) ice-cold artificial cerebrospinal fluid (aCSF) solution: NaCl 124 mM; KCl 3 mM; NaH_2_PO_4_ 1.25 mM; NaHCO_3_ 26 mM; MgSO_4_ 1 mM; CaCl_2_ 2 mM; and glucose 10 mM (pH = 7.4). Slices (400 μm thick) were cut perpendicularly to the long axis of the hippocampus with a McIIwain tissue chopper and allowed to recover for at least 1h in a resting chamber, filled with aCSF, at room temperature. Slices were then transferred to a submerging chamber (1 ml) and continuously superfused with oxigenated aCSF at 32 °C at a flow rate of ~3 mL/min. Evoked field potentials were recorded in the outer molecular layer of the DG, close to the hippocampal fissure, using glass microelectrodes filled with aCSF (2–8 MΩ resistance, Harvard apparatus Ltd., Cambridge, MA, USA) (**Fig. S2f**). Briefly, the lateral path (LPP) was stimulated using an isolated stimulation unit and a concentric bipolar electrode, and test pulses were delivered every 20 seconds for 15 min for obtaining a baseline recording. Due to the small size of field excitatory postsynaptic potentials (fEPSPs), the intensity of stimulus (80–200 μA) was initially adjusted to obtain a large fEPSP slope with minimum population spike contamination. Only recordings displaying paired-pulse depression (50 ms interpulse interval) were accepted. Test pulses were delivered every 20 seconds for 15 minutes for obtaining a baseline recording and LTP was induced using 1 second 100 Hz trains of stimulation, repeated 4 times with a 15 seconds interval. Subsequently, responses were registered every 20 seconds for up to 60 min following the high-frequency stimulation (HFS). Recordings were obtained with an Axoclamp 2B amplifier (Axon Instruments, Foster City, CA, USA), digitized and continuously stored on a computer with the WinLTP software, filtered at 2.9 kHz and sampled at 10–20 kHz. Normalized fEPSP slopes were used to calculate changes in synaptic efficacy following HFS, which were evaluated between 50 and 60 min after HFS.

**Data analysis and statistics**

For all experiments, observations outside of the interval defined by the first quartile (Q1) - 1.5 interquartile range (IQR) and the third quartile (Q3) + 1.5 IQR, were considered outliers and removed from analysis. Given that all data followed a Gaussian distribution, parametric tests were applied. Importantly, although n = 10 per group was firstly established, a threshold (T) based on molecular and behavioural outputs was defined for posterior analyses given the high variability of data (with resilient and susceptible populations being observed in uCMS groups): T = CORT / EPM % open arms, where “CORT” represents the corticosterone raw values and “EPM % open arms” the percentage of time in open arms during EPM test. In all analyses, uCMS animals with values below this defined threshold (specifically, with values below the first quartile (Q1) of uCMS animals), i.e., with similar values as CTRL animals, were excluded from further analysis (Fig. S2e). For summary radar plots (**Fig. 5**), CTRL was set to 100% and all other conditions were normalized to CTRL; the following parameters were used: (a) percentage of time in open arms during EPM, percentage of time spent in immobility behaviour in FST (inverted as a function of CTRL values), percentage of time spent in the centre area in OF test, percentage of time spent exploring the novel object in NOR and latency to groom in SST; (b) total number of Ki67+, DCX+, Ki67+DCX+ and BrdU+NeuN+ cells per total volume (mm^3^); and (c) Iba1, GFAP and MBP occupancy per DG granular area. Data were analysed through two-tailed unpaired and paired student’s t-tests, two-way or three-way analysis of variance (ANOVA), with Dunnett’s correction for multiple comparisons when appropriate (unless stated otherwise). The analyses comparing CTRL *versus* uCMS were performed through student’s t-tests; analyses between the different uCMS groups were performed through two-way or three-way ANOVA depending on the number of variables. Statistical analysis was performed using Prism v.8.4.2 (GraphPad Software, USA). Results are expressed as mean ± standard error of mean (SEM) and statistical significance was set when *p*<0.05.

**Legends to Supplementary Figures and Tables**

**Fig. S1 CB2R ligands dose-response effects on behaviour and AHN.** **a** Schematic representation of the experimental timeline: 2 weeks of pharmacological treatments in physiological conditions followed by behavioural testing. **b** HU308 treatment does not interfere with anxiety-like behaviour (EPM performance). **c** HU308 treatment does not interfere with coping behaviour (FST performance). **d** AM630 treatment at 5 mg/kg dose increases anxiety-like behaviour (decrease in time spent in open arms during EPM). **e** AM630 treatment impairs ability to cope with an inescapable stressor (coping behaviour) at 0.5 mg/kg dose (increase in immobile behaviour during FST). **f** HU308 treatment does not interfere with cognitive performance during NOR test. **g** AM630 treatment (in both doses) impairs ability to recognize novel object during NOR test. **h** HU308 treatment accelerates neurogenesis by increasing the total number of DCX+ (h1) and BrdU+/DCX+ (h2) cells in the hippocampal DG. **i** AM630 treatment does not alter neurogenesis in terms of DCX+ (i1) or BrdU+/DCX+ (i2) cells in the hippocampal DG. **j** Representative images of DG coronal sections of all tested conditions, stained for BrdU (green), DCX (red) and DAPI (blue); scale bar = 50 µm. Data presented as mean ± SEM and circles individual data points (animals) [Student’s t-tests and One-way ANOVA: ****p*<0.001, **p*<0.05 vs CTRL or in familiar (f) vs novel (n) (F,G); further statistical details in **Table S4**]. HU308, CB2R agonist; AM630, CB2R inverse agonist.

**Fig. S2 Experimental outline and physical, biochemical and molecular changes.** **a** Schematic representation of the experimental timeline: 8 weeks of uCMS followed by 2 weeks of mild uCMS and interventional procedures (PE and/or CB2R ligands) and behaviour assessment; four weeks before sacrifice, BrdU (2×50 mg/kg/day) was administered for five consecutive days. **b** uCMS schedule: blood collection was performed at the end of uCMS protocol (week 8). **c** Weight variation (as a percentage of weight change relative to initial weight) of all experimental groups, (c1) CTRL vs all uCMS, (c2) CTRL vs uCMS vs uCMS+PE, (c3) CTRL vs uCMS vs uCMS+HU vs uCMS+HU+PE, (c4) CTRL vs uCMS vs uCMS+AM vs uCMS+AM+PE, during experimental timeline (light gray shading represents interventional period). **d** Corticosterone levels at the end of uCMS protocol, (d1) CTRL vs all uCMS and (d2) variation between all experimental groups. **e** Defined threshold (T = CORT / EPM % open arms) for further analysis (see section 4.7, *Data analysis and statistics*), (e1) CTRL vs all uCMS and (e2) variation between all experimental groups (gray shading represents values below the defined threshold). **f** Schematic representation of the stimulation and recording electrodes in the DG for recording fEPSPs of LPP synapses. uCMS decreases the fEPSP slope of DG slices, which remains unchanged after PE in combination with HU308 or AM630 treatment. **g** CB2R protein levels are tendentially decreased by chronic stress but remain unaltered after PE and treatment with HU308 and/or AM630. Representative bands of CB2R (38 kDa) and GAPDH (37 kDa) expression of all tested conditions and correspond to each bar in the plot. Data presented as mean ± SEM and circles individual data points (animals). [Student’s t-test: ***p*<0.01; further statistical details in **Table S4**]. uCMS, unpredictable chronic mild stress (all uCMS, all uCMS groups); AM630, CB2R inverse agonist; HU308, CB2R agonist; PE, physical exercise.

**Fig. S3 (related to Fig. 1) CB2R activation does not rescue chronic stress-induced behavioural impairments.** **a** Schematic representation of the experimental design. **a** uCMS induces anxiety-like behaviour (decrease in time spent in open arms during EPM), which persists upon HU308 treatment and PE. **c** uCMS-evoked increased latency to groom (in SST) is not altered by HU308 treatment and PE. **d** uCMS-induced immobile behaviour in FST persists with HU308 treatment and PE. **e** No changes in distance travelled in centre zone in the OF test. **f** PE, but not HU308 treatment, rescues uCMS-induced cognitive impairments in NOR. Data presented as mean ± SEM and circles individual data points (animals). [Student’s t-test and Two-way ANOVA; *****p*<0.0001, ***p*<0.01; further statistical details in **Table S4**]. uCMS, unpredictable chronic mild stress; HU308, CB2R agonist; PE, physical exercise.

**Fig. S4 (related to Fig. 4) PE, but not CB2R activation, partially ameliorates chronic stress-induced neuroinflammatory changes. a** uCMS increases Iba1 granular expression, an effect prevented by PE and accompanied with HU308 treatment. **b** PE and/or HU308 treatment do not rescue chronic stress-induced decrease in MBP granular expression. **c** uCMS and HU308 treatment in combination with PE have no effect in GFAP granular expression. The expression of neuroinflammation-related markers **d** TNFα, **e** IL1β, **f** IL10, **g** arginase and **h** iNOS is differentially affected by uCMS and further accentuated by HU308 treatment. Data presented as mean ± SEM and circles individual data points (animals). [Student’s t-tests: *****p*<0.0001, **p*<0.05 CTRL vs uCMS; Two-way ANOVA: ^#^*p*<0.05 vs uCMS; further statistical details in **Table S4**]. uCMS, unpredictable chronic mild stress; HU308, CB2R agonist; PE, physical exercise.

**Table S1 – Key resources table**

**Table S2 – Estimated DG volumes (mm^3^) in tested experimental conditions (data as mean ± SEM)**

**Table S3 – List of primers**

**Table S4 – Statistical summary of results**

**Supplementary Tables**

**Table S1 – Key resources table**

| **Reagent or Resource** | **Source(s)** | **Identifier(s)** | **Other information** |
| --- | --- | --- | --- |
| ***Antibodies*** | | | |
| Anti-BrdU (mouse monoclonal) | Dako, Agilent Technologies | #M0744, RRID:AB_10013660 | IF (1:500) |
| Anti-BrdU (rat monoclonal) | Abcam | #ab6326, RRID:AB_305426 | IF (1:500) |
| Anti-DCX (chicken polyclonal) | Synaptic Systems | #326 006, RRID:AB_2737040 | IF (1:750) |
| Anti- DCX (goat polyclonal) | Santa Cruz Biotechnologies | #sc-8066, RRID:AB_2088494 | IF (1:500) |
| Anti-GFAP (mouse monoclonal) | Novocastra, Leica Biosystems | #NCL-GFAP-GA5, RRID:AB_563739 | IF (1:100) |
| Anti-Iba1 (rabbit polyclonal) | FUJIFILM Wako Chemicals Europe GmbH | #010-19741 | IF (1:250) |
| Anti-Ki67 (rabbit monoclonal) | Abcam | #ab16667, RRID:AB_302459 | IF (1:500) |
| Anti-MBP (rat monoclonal) | Bio-Rad Laboratories | #MCA409S, RRID:AB_325004 | IF (1:200) |
| Anti-NeuN (rabbit monoclonal) | Cell Signalling Technology | #12943, RRID:AB_2630395 | IF (1:500) |
| Alexa Fluor 488 (goat anti-mouse polyclonal) | Thermo Fisher Scientific | #A-11001, RRID:AB_2534069 | IF (1:500) |
| Alexa Fluor 488 (goat anti-chicken polyclonal) | Thermo Fisher Scientific | #A-11039, RRID:AB_2534096 | IF (1:1000) |
| Alexa Fluor 488 (goat anti-rabbit polyclonal) | Thermo Fisher Scientific | #A-11008, RRID:AB_143165 | IF (1:500) |
| Alexa Fluor 488 (goat anti-rat polyclonal) | Thermo Fisher Scientific | #A-11006, RRID:AB_2534074 | IF (1:1000) |
| Alexa Fluor 568 (goat anti-goat polyclonal) | Thermo Fisher Scientific | #A-11057, RRID:AB_2534104 | IF (1:500) |
| Alexa Fluor 568 (goat anti-rabbit polyclonal) | Thermo Fisher Scientific | #A-11011, RRID:AB_143157 | IF (1:1000) |
| Alexa Fluor 568 (goat anti-rat polyclonal) | Thermo Fisher Scientific | #A-11077, RRID:AB_2534121 | IF (1:500) |
| Alexa Fluor 647 (goat anti-mouse polyclonal) | Thermo Fisher Scientific | #A-21235, RRID:AB_2535804 | IF (1:500) |
| Anti-CB1R (mouse monoclonal) | Santa Cruz Biotechnologies | #sc-293419 | WB (1:1000) |
| Anti-CB2R (mouse monoclonal) | Santa Cruz Biotechnologies | sc-293188 | WB (1:1000) |
| Anti-GAPDH (mouse monoclonal) | Thermo Fisher Scientific | #AM4300, RRID:AB_2536381 | WB (1:5000) |
| Goat anti-mouse IgG (H+L)-HRP conjugate | Bio-Rad Laboratories | #172-1011, RRID:AB_11125936 | WB (1:5000) |
| ***Chemicals, peptides and proteins*** | | | |
| HU308 (Cannabinoid CB2 receptor selective agonist) | Tocris | #3088 | “HU” for short |
| AM630 (Cannabinoid CB2 receptor selective inverse agonist) |  | #1120 | “AM” for short |
| BrdU (5-Bromo-2’deoxyuridine) | Sigma Aldrich | #B5002 |  |
| DAPI (4',6-diamidino-2-phenylindole) | Sigma Aldrich | #D9542 |  |
| ***Commercial assays*** | | | |
| Corticosterone ELISA Kit | Abcam | #ab108821, RRID:AB_2889904 |  |
| ***Experimental models: organisms/strains*** | | | |
| Mouse (*Mus musculus,* C57BL/6J) | Charles River Laboratories |  | Male mice  (14 weeks-old) |
| ***Equipment*** | | | |
| Treadmill apparatus  (LE8710MTS model) | Panlab, Harvard Apparatus | #76-0896 |  |
| ***Software and algorithms*** | | | |
| Fiji software | Open-source software | RRID:SCR_002285 | <https://imagej.net/software/fiji/> |
| Prism v.8.4.2 | GraphPad Software Inc | RRID:SCR_002798 |  |
| SMART® v2.5 | Panlab, Harvard Apparatus | RRID:SCR_002852 |  |
| Solomon Coder beta 17.03.22 | Open-source software | RRID:SCR_016041 | <https://solomon.andraspeter.com/> |
| WinLTP software | WinLTP Ltd. | RRID:SCR_008590 |  |

| **Table S2 – Estimated DG volumes (mm^3^) in tested experimental conditions (data as mean ± SEM)** | | | | | |
| --- | --- | --- | --- | --- | --- |
|  | **CTRL** | **uCMS** | **uCMS+PE** | **uCMS+PE+AM** | **uCMS+AM** |
| Total | 4.7 (±0.4) x10^-2^ | 4.2 (±0.2) x10^-2^ | 4.3 (±0.2) x10^-2^ | 4.7 (±0.1) x10^-2^ | 4.1 (±0.1) x10^-2^ |
| Dorsal | 1.5 (±0.1) x10^-2^ | 1.2 (±0.06) x10^-2^ | 1.7 (±0.08) x10^-2^ | 1.5 (±0.4) x10^-2^ | 1.4 (±0.1) x10^-2^ |
| Ventral | 3.1 (±0.7) x10^-2^ | 2.9 (±0.6) x10^-2^ | 2.6 (±0.3) x10^-2^ | 2.6 (±0.1) x10^-2^ | 2.6 (±0.5) x10^-2^ |

| **Table S3 – List of primers** | | |
| --- | --- | --- |
| **Gene** | **Forward primer sequence** | **Reverse primer sequence** |
| TNF-α | 5’-TACTGAACTTCGGGGTGATTGGTCC-3’ | 5’-CAGCCTTGTCCCTTGAAGAGAACC-3’ |
| IL-1β | 5'- CAGGCTCCGAGATGAACAAC-3' | 5'- GGTGGAGAGCTTTCAGCTCATA-3' |
| IL-10 | 5’-CCAGTTTTACCTGGTAGAAGTGAG-3’ | 5’-TGTCTAGGTCCTGGAGTCCAGCAGACTC-3’ |
| Arginase 1 | 5’-CTTGGCTTGCTTCGGAACTC-3’ | 5’-GGAGAAGGCGTTTGCTTAGTTC-3’ |
| iNOS | 5’-ACCCACATCTGGCAGAATGAG-3’ | 5’-AGCCATGACCTTTCGCATTAG-3’ |
| β-actin | 5’-GCTCCGGCATGTGCAA-3’ | 5’-AGGATCTTCATGAGGTAGT-3’ |

| **Table S4 – Statistical summary of results** | | | | | |
| --- | --- | --- | --- | --- | --- |
| **Figure** | **Experiment** | **Statistics** | **Comparisons** | **Statistical details** | |
|  |  |  |  | **t / F** | ***p* value** |
| **Figure 1b** | EPM (% time open arms) | Student’s  t-test | CTRL vs uCMS (unpaired) | t_16_ = 3.492 | ***0.0030***** |
|  |  | Two-way ANOVA | AM | F (1, 27) = 3.889 | 0.0589 |
|  |  |  | PE | F (1, 27) = 5.982 | ***0.0213**** |
|  |  |  | PE*AM | F (1, 27) = 0.2909 | 0.5941 |
|  |  | Post-hoc | uCMS vs uCMS+PE |  | 0.4059 |
|  |  |  | uCMS vs uCMS+AM+PE |  | ***0.0079***** |
|  |  |  | uCMS vs uCMS+AM |  | 0.6236 |
| **Figure 1c** | SST (latency) | Student’s  t-test | CTRL vs uCMS (unpaired) | t_16_ = 3.941 | ***0.0012***** |
|  |  | Two-way ANOVA | AM | F (1, 27) = 2.473 | 0.1275 |
|  |  |  | PE | F (1, 27) = 6.764 | ***0.0149**** |
|  |  |  | PE*AM | F (1, 27) = 0.5974 | 0.04463 |
|  |  | Post-hoc | uCMS vs uCMS+PE |  | 0.4904 |
|  |  |  | uCMS vs uCMS+AM+PE |  | ***0.0133**** |
|  |  |  | uCMS vs uCMS+AM |  | 0.9212 |
| **Figure 1d** | FST (time in immobile behaviour) | Student’s  t-test | CTRL vs uCMS (unpaired) | t_16_ = 6.087 | ***<0.0001 ******* |
|  |  | Two-way ANOVA | AM | F (1, 27) = 9.930 | ***0.0040***** |
|  |  |  | PE | F (1, 27) = 1.950 | 0.1739 |
|  |  |  | PE*AM | F (1, 27) = 0.1998 | 0.6585 |
|  |  | Post-hoc | uCMS vs uCMS+PE |  | 0.4326 |
|  |  |  | uCMS vs uCMS+AM+PE |  | ***0.0062***** |
|  |  |  | uCMS vs uCMS+AM |  | ***0.0419**** |
| **Figure 1e** | OF (% time in centre) | Student’s  t-test | CTRL vs uCMS (unpaired) | t_15_ = 1.800 | 0.0920 |
|  |  | Two-way ANOVA | AM | F (1, 26) = 0.4285 | 0.5185 |
|  |  |  | PE | F (1, 26) = 3.275 | 0.0819 |
|  |  |  | PE*AM | F (1, 26) = 0.1820 | 0.1889 |
| **Figure 1f** | NOR (% exploration time) | Student’s  t-tests (paired) | CTRL_f vs CTRL_n | t_8_ = 8.525 | ***<0.0001 ******* |
|  |  |  | uCMS_f vs uCMS_n | t_8_ = 0.8505 | 0.4198 |
|  |  |  | uCMS+PE_f vs uCMS+PE_n | t_6_ = 4.445 | ***0.0044***** |
|  |  |  | uCMS+AM+PE_f vs uCMS+AM+PE_n | t_7_ = 5.473 | ***0.0009****** |
|  |  |  | uCMS+AM_f vs uCMS+AM_n | t_6_ = 2.678 | ***0.0366**** |
| **Figure 2a1** | Cell proliferation (total Ki67+ cells) | Student’s t-test | CTRL vs uCMS (unpaired) | t_8_ = 6.564 | ***0.0002****** |
|  |  | Two-way ANOVA | AM | F (1, 13) = 12.81 | ***0.0034***** |
|  |  |  | PE | F (1, 13) = 36.87 | ***<0.0001 ******* |
|  |  |  | PE*AM | F (1, 13) = 3.698 | 0.0767 |
|  |  | Post-hoc | uCMS vs uCMS+PE |  | ***0.0263**** |
|  |  |  | uCMS vs uCMS+AM+PE |  | ***<0.00001 ******* |
|  |  |  | uCMS vs uCMS+AM |  | 0.5184 |
| **Figure 2a2** | Cell proliferation (dorsal Ki67+ cells) | Student’s  t-test | CTRL vs uCMS (unpaired) | t_8_ = 2.298 | 0.00506 |
|  |  | Two-way ANOVA | AM | F (1, 13) = 0.1256 | 0.7287 |
|  |  |  | PE | F (1, 13) = 9.519 | ***0.0087***** |
|  |  |  | PE*AM | F (1, 13) = 3.354 | 0.0901 |
|  |  | Post-hoc | uCMS vs uCMS+PE |  | 0.7061 |
|  |  |  | uCMS vs uCMS+AM+PE |  | 0.0681 |
|  |  |  | uCMS vs uCMS+AM |  | 0.6009 |
| **Figure 2a3** | Cell proliferation (ventral Ki67+ cells) | Student’s  t-test | CTRL vs uCMS (unpaired) | t_8_ = 6.236 | ***0.0002****** |
|  |  | Two-way ANOVA | AM | F (1, 13) = 7.527 | ***0.0167**** |
|  |  |  | PE | F (1, 13) = 12.07 | ***0.0041***** |
|  |  |  | PE*AM | F (1, 13) = 1.761 | 0.2074 |
|  |  | Post-hoc | uCMS vs uCMS+PE |  | 0.3224 |
|  |  |  | uCMS vs uCMS+AM+PE |  | 0.6294 |
|  |  |  | uCMS vs uCMS+AM |  | ***0.0016***** |
| **Figure 2b1** | Neuronal differentiation (total DCX+ cells) | Student’s  t-test | CTRL vs uCMS (unpaired) | t_8_ = 5.232 | ***0.0008****** |
|  |  | Two-way ANOVA | AM | F (1, 13) = 0.6126 | 0.4478 |
|  |  |  | PE | F (1, 13) = 18.74 | ***0.0008****** |
|  |  |  | PE*AM | F (1, 13) = 0.1552 | 0.7000 |
|  |  | Post-hoc | uCMS vs uCMS+PE |  | ***0.0352**** |
|  |  |  | uCMS vs uCMS+AM+PE |  | ***0.0071***** |
|  |  |  | uCMS vs uCMS+AM |  | 0.9852 |
| **Figure 2b2** | Neuronal differentiation (dorsal DCX+ cells) | Student’s  t-test | CTRL vs uCMS (unpaired) | t_8_ = 4.375 | ***0.0024***** |
|  |  | Two-way ANOVA | AM | F (1, 13) = 1.929 | 0.1882 |
|  |  |  | PE | F (1, 13) = 10.50 | ***0.0064***** |
|  |  |  | PE*AM | F (1, 13) = 0.2425 | 0.6306 |
|  |  | Post-hoc | uCMS vs uCMS+PE |  | 0.1630 |
|  |  |  | uCMS vs uCMS+AM+PE |  | 0.4337 |
|  |  |  | uCMS vs uCMS+AM |  | 0.4218 |
| **Figure 2b3** | Neuronal differentiation (ventral DCX+ cells) | Student’s  t-test | CTRL vs uCMS (unpaired) | t_8_ = 3.667 | ***0.0063***** |
|  |  | Two-way ANOVA | AM | F (1, 13) = 1.184 | 0.2963 |
|  |  |  | PE | F (1, 13) = 8.151 | ***0.0135**** |
|  |  |  | PE*AM | F (1, 13) = 0.4409 | 0.5183 |
|  |  | Post-hoc | uCMS vs uCMS+PE |  | 0.0613 |
|  |  |  | uCMS vs uCMS+AM+PE |  | ***0.0348**** |
|  |  |  | uCMS vs uCMS+AM |  | 0.4758 |
| **Figure 2c1** | Neuronal “acceleration” (total Ki76+DCX+ cells) | Student’s  t-test | CTRL vs uCMS (unpaired) | t_8_ = 10.46 | ***<0.0001 ******* |
|  |  | Two-way ANOVA | AM | F (1, 13) = 5.715 | ***0.0326**** |
|  |  |  | PE | F (1, 13) = 15.89 | ***0.0016***** |
|  |  |  | PE*AM | F (1, 13) = 0.3185 | 0.5821 |
|  |  | Post-hoc | uCMS vs uCMS+PE |  | 0.0697 |
|  |  |  | uCMS vs uCMS+AM+PE |  | ***0.0013***** |
|  |  |  | uCMS vs uCMS+AM |  | 0.4443 |
| **Figure 2c2** | Neuronal “acceleration” (dorsal Ki76+DCX+ cells) | Student’s  t-test | CTRL vs uCMS (unpaired) | t_8_ = 4.905 | ***0.0012***** |
|  |  | Two-way ANOVA | AM | F (1, 13) = 0.0263 | 0.8737 |
|  |  |  | PE | F (1, 13) = 4.348 | 0.0573 |
|  |  |  | PE*AM | F (1, 13) = 8.923 | ***0.0105**** |
|  |  | Post-hoc | uCMS vs uCMS+PE |  | 0.8579 |
|  |  |  | uCMS vs uCMS+AM+PE |  | 0.2898 |
|  |  |  | uCMS vs uCMS+AM |  | 0.1485 |
| **Figure 2c3** | Neuronal “acceleration” (ventral Ki76+DCX+ cells) | Student’s  t-test | CTRL vs uCMS (unpaired) | t_8_ = 4.442 | ***0.0022***** |
|  |  | Two-way ANOVA | AM | F (1, 8) = 5.682 | ***0.0443**** |
|  |  |  | PE | F (1, 8) = 3.622 | 0.1706 |
|  |  |  | PE*AM | F (1, 8) = 0.06082 | 0.8061 |
|  |  | Post-hoc | uCMS vs uCMS+PE |  | 0.3203 |
|  |  |  | uCMS vs uCMS+AM+PE |  | ***0.0322**** |
|  |  |  | uCMS vs uCMS+AM |  | 0.2596 |
| **Figure 2e1** | Neuronal survival (total BrdU+NeuN+ cells) | Student’s  t-test | CTRL vs uCMS (unpaired) | t_8_ = 7.314 | ***<0.0001 ******* |
|  |  | Two-way ANOVA | AM | F (1, 13) = 3.928 | 0.0690 |
|  |  |  | PE | F (1, 13) = 14.47 | ***0.0022***** |
|  |  |  | PE*AM | F (1, 13) = 0.3412 | 0.5691 |
|  |  | Post-hoc | uCMS vs uCMS+PE |  | 0.0905 |
|  |  |  | uCMS vs uCMS+AM+PE |  | ***0.0028***** |
|  |  |  | uCMS vs uCMS+AM |  | 0.6383 |
| **Figure 2e2** | Neuronal survival (dorsal BrdU+NeuN+ cells) | Student’s  t-test | CTRL vs uCMS (unpaired) | t_8_ = 1.877 | 0.0973 |
|  |  | Two-way ANOVA | AM | F (1, 13) = 0.7715 | 0.3957 |
|  |  |  | PE | F (1, 13) = 0.1311 | 0.7231 |
|  |  |  | PE*AM | F (1, 13) = 1.016 | 0.3318 |
| **Figure 2e3** | Neuronal survival (ventral BrdU+NeuN+ cells) | Student’s  t-test | CTRL vs uCMS (unpaired) | t_8_ = 3.786 | ***0.0053***** |
|  |  | Two-way ANOVA | AM | F (1, 13) = 3.225 | 0.0958 |
|  |  |  | PE | F (1, 13) = 25.46 | ***0.0002****** |
|  |  |  | PE*AM | F (1, 13) = 3.328 | 0.0912 |
|  |  | Post-hoc | uCMS vs uCMS+PE |  | 0.1039 |
|  |  |  | uCMS vs uCMS+AM+PE |  | ***0.0008****** |
|  |  |  | uCMS vs uCMS+AM |  | 0.9993 |
| **Figure 3b1** | Cell proliferation SGZ/GCL subregional preference (dorsal Ki67+ cells) | Student’s  t-tests (paired) | CTRL_supra: SGZ vs GCL | t_4_ = 12.79 | ***0.0009****** |
|  |  |  | CTRL_infra: SGZ vs GCL | t_4_ = 9.20 | ***0.0027***** |
|  |  |  | uCMS_supra: SGZ vs GCL | t_4_ = 4.846 | ***0.0280**** |
|  |  |  | uCMS_infra: SGZ vs GCL | t_4_ = 9.86 | ***0.0035***** |
|  |  |  | uCMS+PE_supra: SGZ vs GCL | t_3_ = 8.766 | ***0.0031***** |
|  |  |  | uCMS+PE_infra: SGZ vs GCL | t_3_ = 12.29 | ***0.0026***** |
|  |  |  | uCMS+AM+PE_supra: SGZ vs GCL | t_3_ = 12.55 | ***0.0021***** |
|  |  |  | uCMS+AM+PE_infra: SGZ vs GCL | t_3_ = 6.652 | ***0.0219**** |
|  |  |  | uCMS+AM_supra: SGZ vs GCL | t_3_ = 12.71 | ***0.0061***** |
|  |  |  | uCMS+AM_infra: SGZ vs GCL | t_3_ = 3.380 | 0.0775 |
| **Figure 3b2** | Cell proliferation SGZ/GCL subregional preference (ventral Ki67+ cells) | Student’s  t-tests (paired) | CTRL_supra: SGZ vs GCL | t_4_ = 12.347 | ***0.00012****** |
|  |  |  | CTRL_infra: SGZ vs GCL | t_4_ = 8.054 | ***0.0013***** |
|  |  |  | uCMS_supra: SGZ vs GCL | t_4_ = 3.346 | ***0.0287**** |
|  |  |  | uCMS_infra: SGZ vs GCL | t_4_ = 0.8065 | 0.4651 |
|  |  |  | uCMS+PE_supra: SGZ vs GCL | t_3_ = 5.097 | ***0.0146**** |
|  |  |  | uCMS+PE_infra: SGZ vs GCL | t_3_ = 5.450 | ***0.0121**** |
|  |  |  | uCMS+AM+PE_supra: SGZ vs GCL | t_3_ = 15.78 | ***0.0040***** |
|  |  |  | uCMS+AM+PE_infra: SGZ vs GCL | t_3_ = 20.31 | ***0.0024***** |
|  |  |  | uCMS+AM_supra: SGZ vs GCL | t_3_ = 1.598 | 0.2083 |
|  |  |  | uCMS+AM_infra: SGZ vs GCL | t_3_ = 2.957 | 0.0597 |
| **Figure 3c1** | Cell proliferation supra/infra subregional preference (dorsal Ki67+ cells) | Student’s  t-tests (paired) | CTRL_SGZ: supra vs infra | t_4_ = 5.693 | ***0.0047***** |
|  |  |  | CTRL_GCL: supra vs infra | t_4_ = 2.158 | 0.0971 |
|  |  |  | uCMS_SGZ: supra vs infra | t_4_ = 0.1747 | 0.8698 |
|  |  |  | uCMS_GCL: supra vs infra | t_4_ = 2.949 | ***0.0420**** |
|  |  |  | uCMS+PE_SGZ: supra vs infra | t_3_ = 1.857 | 0.1604 |
|  |  |  | uCMS+PE_GCL: supra vs infra | t_3_ = 0.3016 | 0.7826 |
|  |  |  | uCMS+AM+PE_SGZ: supra vs infra | t_3_ = 3.985 | ***0.0283**** |
|  |  |  | uCMS+AM+PE_GCL: supra vs infra | t_3_ = 1.000 | 0.3910 |
|  |  |  | uCMS+AM_SGZ: supra vs infra | t_3_ = 0.8960 | 0.4363 |
|  |  |  | uCMS+AM_GCL: supra vs infra | t_3_ = 2.259 | 0.1091 |
| **Figure 3c2** | Cell proliferation supra/infra subregional preference (ventral Ki67+ cells) | Student’s  t-tests (paired) | CTRL_SGZ: supra vs infra | t_4_ = 2.978 | ***0.0408**** |
|  |  |  | CTRL_GCL: supra vs infra | t_4_ = 2.825 | ***0.0476**** |
|  |  |  | uCMS_SGZ: supra vs infra | t_4_ = 1.833 | 0.1408 |
|  |  |  | uCMS_GCL: supra vs infra | t_4_ = 2.672 | 0.0557 |
|  |  |  | uCMS+PE_SGZ: supra vs infra | t_3_ = 1.255 | 0.2982 |
|  |  |  | uCMS+PE_GCL: supra vs infra | t_3_ = 1.260 | 0.2967 |
|  |  |  | uCMS+AM+PE_SGZ: supra vs infra | t_3_ = 9.706 | ***0.0023***** |
|  |  |  | uCMS+AM+PE_GCL: supra vs infra | t_3_ = 3.446 | ***0.0411**** |
|  |  |  | uCMS+AM_SGZ: supra vs infra | t_3_ = 0.7395 | 0.5132 |
|  |  |  | uCMS+AM_GCL: supra vs infra | t_3_ = 0.9227 | 0.4242 |
| **Figure 3d1** | Neuronal differentiation (DCX+ sub populations) | Student’s  t-tests (paired) | CTRL: progenitor vs immature | t_4_ = 20.68 | ***<0.0001 ******* |
|  |  |  | uCMS: progenitor vs immature | t_4_ = 7.498 | ***0.0017***** |
|  |  |  | uCMS+PE: progenitor vs immature | t_3_ = 5.719 | ***0.0106**** |
|  |  |  | uCMS+AM+PE: progenitor vs immature | t_3_ = 8.979 | ***0.0029***** |
|  |  |  | uCMS+AM: progenitor vs immature | t_3_ = 5.825 | ***0.0101**** |
| **Figure 3d2** | Neuronal differentiation dorsal/ventral subregional preference (DCX+ sub populations) | Student’s  t-tests (paired) | CTRL_dorsal: progenitor vs immature | t_4_ = 22.98 | ***<0.0001 ******* |
|  |  |  | CTRL_ventral: progenitor vs immature | t_4_ = 21.00 | ***<0.0001 ******* |
|  |  |  | uCMS_dorsal: progenitor vs immature | t_4_ = 6.302 | ***0.0032***** |
|  |  |  | uCMS_ventral: progenitor vs immature | t_4_ = 5.917 | ***0.0041***** |
|  |  |  | uCMS+PE_dorsal: progenitor vs immature | t_3_ = 13.34 | ***0.0009****** |
|  |  |  | uCMS+PE_ventral: progenitor vs immature | t_3_ = 10.30 | ***0.0020***** |
|  |  |  | uCMS+AM+PE_dorsal: progenitor vs immature | t_3_ = 13.57 | ***0.0009****** |
|  |  |  | uCMS+AM+PE_ventral: progenitor vs immature | t_3_ = 9.625 | ***0.0024***** |
|  |  |  | uCMS+AM_dorsal: progenitor vs immature | t_3_ = 4.498 | ***0.0205**** |
|  |  |  | uCMS+AM_ventral: progenitor vs immature | t_3_ = 8.174 | ***0.0038***** |
| **Figure 3d3** | Neuronal differentiation supra/ventral subregional preference (DCX+ sub populations) | Student’s  t-tests (paired) | CTRL_supra: progenitor vs immature | t_4_ = 25.30 | ***<0.0001 ******* |
|  |  |  | CTRL_infra: progenitor vs immature | t_4_ = 7.663 | ***0.0016***** |
|  |  |  | uCMS_supra: progenitor vs immature | t_4_ = 5.227 | ***0.0064***** |
|  |  |  | uCMS_infra: progenitor vs immature | t_4_ = 7.158 | ***0.0020***** |
|  |  |  | uCMS+PE_supra: progenitor vs immature | t_3_ = 9.712 | ***0.0023***** |
|  |  |  | uCMS+PE_infra: progenitor vs immature | t_3_ = 14.01 | ***0.0008****** |
|  |  |  | uCMS+AM+PE_supra: progenitor vs immature | t_3_ = 8.139 | ***0.0039***** |
|  |  |  | uCMS+AM+PE_infra: progenitor vs immature | t_3_ = 12.92 | ***0.0010***** |
|  |  |  | uCMS+AM_supra: progenitor vs immature | t_3_ = 2.999 | 0.0577 |
|  |  |  | uCMS+AM_infra: progenitor vs immature | t_3_ = 4.375 | ***0.0279**** |
| **Figure 3e1** | Neuronal “acceleration” dorsal/ventral subregional preference (Ki76+DCX+ cells) | Student’s  t-tests (paired) | CTRL: dorsal vs ventral | t_4_ = 1.473 | 0.2147 |
|  |  |  | uCMS: dorsal vs ventral | t_4_ = 4.084 | ***0.0151**** |
|  |  |  | uCMS+PE: dorsal vs ventral | t_3_ = 1.956 | 0.1454 |
|  |  |  | uCMS+AM+PE: dorsal vs ventral | t_3_ = 1.599 | 0.2082 |
|  |  |  | uCMS+AM: dorsal vs ventral | t_3_ = 0.5055 | 0.6480 |
| **Figure 3e2** | Neuronal “acceleration” supra/infra subregional preference (Ki76+DCX+ cells) | Student’s  t-tests (paired) | CTRL: supra vs infra | t_4_ = 6.185 | ***0.0035***** |
|  |  |  | uCMS: supra vs infra | t_4_ = 0.1943 | 0.8554 |
|  |  |  | uCMS+PE: supra vs infra | t_3_ = 1.238 | 0.3038 |
|  |  |  | uCMS+AM+PE: supra vs infra | t_3_ = 5.255 | ***0.0134**** |
|  |  |  | uCMS+AM: supra vs infra | t_3_ = 2.174 | 0.1180 |
| **Figure 3g1** | Neuronal survival SGZ/GCL subregional preference (dorsal BrdU+NeuN+ cells) | Student’s  t-tests (paired) | CTRL_supra: SGZ vs GCL | t_4_ = 11.89 | ***0.0003****** |
|  |  |  | CTRL_infra: SGZ vs GCL | t_4_ = 2.033 | 0.1118 |
|  |  |  | uCMS_supra: SGZ vs GCL | t_4_ = 0.7255 | 0.5083 |
|  |  |  | uCMS_infra: SGZ vs GCL | t_4_ = 1.403 | 0.2333 |
|  |  |  | uCMS+PE_supra: SGZ vs GCL | t_3_ = 16.30 | ***0.0005****** |
|  |  |  | uCMS+PE_infra: SGZ vs GCL | t_3_ = 0.7745 | 0.4951 |
|  |  |  | uCMS+AM+PE_supra: SGZ vs GCL | t_3_ = 5.344 | ***0.0128**** |
|  |  |  | uCMS+AM+PE_infra: SGZ vs GCL | t_3_ = 1.567 | 0.2152 |
|  |  |  | uCMS+AM_supra: SGZ vs GCL | t_3_ = 0.9739 | 0.4019 |
|  |  |  | uCMS+AM_infra: SGZ vs GCL | t_3_ = 2.979 | 0.0587 |
| **Figure 3g2** | Neuronal survival SGZ/GCL subregional preference (ventral BrdU+NeuN+ cells) | Student’s  t-tests (paired) | CTRL_supra: SGZ vs GCL | t_4_ = 3.582 | ***0.0231**** |
|  |  |  | CTRL_infra: SGZ vs GCL | t_4_ = 1.965 | 0.1209 |
|  |  |  | uCMS_supra: SGZ vs GCL | t_4_ = 0.3183 | 0.7662 |
|  |  |  | uCMS_infra: SGZ vs GCL | t_4_ = 0.8368 | 0.4498 |
|  |  |  | uCMS+PE_supra: SGZ vs GCL | t_3_ = 2.095 | 0.1271 |
|  |  |  | uCMS+PE_infra: SGZ vs GCL | t_3_ = 1.726 | 0.1829 |
|  |  |  | uCMS+AM+PE_supra: SGZ vs GCL | t_3_ = 6.765 | ***0.0066***** |
|  |  |  | uCMS+AM+PE_infra: SGZ vs GCL | t_3_ = 4.872 | ***0.0165**** |
|  |  |  | uCMS+AM_supra: SGZ vs GCL | t_3_ = 1.769 | 0.1750 |
|  |  |  | uCMS+AM_infra: SGZ vs GCL | t_3_ = 1.000 | 0.3910 |
| **Figure 3h1** | Neuronal survival supra/infra subregional preference (dorsal BrdU+NeuN+ cells) | Student’s  t-tests (paired) | CTRL_SGZ: supra vs infra | t_4_ = 5.235 | ***0.0064***** |
|  |  |  | CTRL_GCL: supra vs infra | t_4_ = 0.3596 | 0.7373 |
|  |  |  | uCMS_SGZ: supra vs infra | t_4_ = 0.09633 | 0.9279 |
|  |  |  | uCMS_GCL: supra vs infra | t_4_ = 0.5345 | 0.6213 |
|  |  |  | uCMS+PE_SGZ: supra vs infra | t_3_ = 3.612 | ***0.0365**** |
|  |  |  | uCMS+PE_GCL: supra vs infra | t_3_ = 1.000 | 0.3910 |
|  |  |  | uCMS+AM+PE_SGZ: supra vs infra | t_3_ = 4.906 | ***0.0162**** |
|  |  |  | uCMS+AM+PE_GCL: supra vs infra | t_3_ = 0.3203 | 0.7698 |
|  |  |  | uCMS+AM_SGZ: supra vs infra | t_3_ = 1.715 | 0.1849 |
|  |  |  | uCMS+AM_GCL: supra vs infra | t_3_ = 0.3203 | 0.7698 |
| **Figure 3h2** | Neuronal survival supra/infra subregional preference (ventral BrdU+NeuN+ cells) | Student’s  t-tests (paired) | CTRL_SGZ: supra vs infra | t_4_ = 6.602 | ***0.0027***** |
|  |  |  | CTRL_GCL: supra vs infra | t_4_ = 0.08114 | 0.9392 |
|  |  |  | uCMS_SGZ: supra vs infra | t_4_ = 1.526 | 0.2016 |
|  |  |  | uCMS_GCL: supra vs infra | t_4_ = 0.3547 | 0.7407 |
|  |  |  | uCMS+PE_SGZ: supra vs infra | t_3_ = 3.470 | ***0.0404**** |
|  |  |  | uCMS+PE_GCL: supra vs infra | t_3_ = 0.1738 | 0.8731 |
|  |  |  | uCMS+AM+PE_SGZ: supra vs infra | t_3_ = 3.283 | ***0.0487**** |
|  |  |  | uCMS+AM+PE_GCL: supra vs infra | t_3_ = 1.997 | 0.1166 |
|  |  |  | uCMS+AM_SGZ: supra vs infra | t_3_ = 0.7144 | 0.5266 |
|  |  |  | uCMS+AM_GCL: supra vs infra | t_3_ = 0.5773 | 0.6042 |
| **Figure 4a** | Iba1 occupancy | Student’s  t-test | CTRL vs uCMS (unpaired) | t_4_ = 2.798 | ***0.0489**** |
|  |  | Two-way ANOVA | AM | F (1, 8) = 9.730 | ***0.0142**** |
|  |  |  | PE | F (1, 8) = 15.92 | ***0.0040***** |
|  |  |  | PE*AM | F (1, 8) = 3.637 | 0.0929 |
|  |  | Post-hoc | uCMS vs uCMS+PE |  | ***0.0080***** |
|  |  |  | uCMS vs uCMS+AM+PE |  | ***0.0026***** |
|  |  |  | uCMS vs uCMS+AM |  | ***0.0187**** |
| **Figure 4b** | MBP occupancy | Student’s  t-test | CTRL vs uCMS (unpaired) | t_4_ = 30.03 | ***<0.0001 ******* |
|  |  | Two-way ANOVA | AM | F (1, 8) = 13.92 | ***0.0058***** |
|  |  |  | PE | F (1, 8) = 1.404 | 0.2701 |
|  |  |  | PE*AM | F (1, 8) = 4.659 | 0.0629 |
|  |  | Post-hoc | uCMS vs uCMS+PE |  | 0.1075 |
|  |  |  | uCMS vs uCMS+AM+PE |  | ***0.0209**** |
|  |  |  | uCMS vs uCMS+AM |  | ***0.0080***** |
| **Figure 4c** | GFAP occupancy | Student’s  t-test | CTRL vs uCMS (unpaired) | t_4_ = 1.302 | 0.2627 |
|  |  | Two-way ANOVA | AM | F (1, 8) = 0.09340 | 0.7677 |
|  |  |  | PE | F (1, 8) = 0.0002040 | 0.9890 |
|  |  |  | PE*AM | F (1, 8) = 0.008523 | 0.7677 |
| **Figure 4e** | TNFα expression | Student’s  t-test | CTRL vs uCMS (unpaired) | t_11_ = 1.456 | 0.1732 |
|  |  | Two-way ANOVA | AM | F (1, 19) = 7.283 | ***0.0142**** |
|  |  |  | PE | F (1, 19) = 8.740 | ***0.0081***** |
|  |  |  | PE*AM | F (1, 19) = 0.7629 | 0.3933 |
|  |  | Post-hoc | uCMS vs uCMS+PE |  | ***0.0317**** |
|  |  |  | uCMS vs uCMS+AM+PE |  | ***0.0017***** |
|  |  |  | uCMS vs uCMS+AM |  | 0.0596 |
| **Figure 4f** | IL-1β expression | Student’s  t-test | CTRL vs uCMS (unpaired) | t_10_ = 2.165 | 0.0557 |
|  |  | Two-way ANOVA | AM | F (1, 20) = 1.211 | 0.2811 |
|  |  |  | PE | F (1, 20) = 3.118 | 0.0927 |
|  |  |  | PE*AM | F (1, 20) = 0.09053 | 0.7666 |
| **Figure 4g** | IL-10 expression | Student’s  t-test | CTRL vs uCMS (unpaired) | t_10_ = 0.7400 | 0.4763 |
|  |  | Two-way ANOVA | AM | F (1, 19) = 1.303 | 0.2678 |
|  |  |  | PE | F (1, 19) = 0.5558 | 0.4651 |
|  |  |  | PE*AM | F (1, 19) = 0.1015 | 0.7535 |
| **Figure 4h** | Arginase expression | Student’s  t-test | CTRL vs uCMS (unpaired) | t_11_ = 1.302 | 0.2196 |
|  |  | Two-way ANOVA | AM | F (1, 20) = 6.504 | ***0.0191**** |
|  |  |  | PE | F (1, 20) = 0.000426 | 0.9837 |
|  |  |  | PE*AM | F (1, 20) = 0.02561 | 0.8745 |
|  |  | Post-hoc | uCMS vs uCMS+PE |  | 0.9985 |
|  |  |  | uCMS vs uCMS+AM+PE |  | 0.1715 |
|  |  |  | uCMS vs uCMS+AM |  | 0.1904 |
| **Figure 4i** | iNOS expression | Student’s  t-test | CTRL vs uCMS (unpaired) | t_11_ = 1.070 | 0.3075 |
|  |  | Two-way ANOVA | AM | F (1, 20) = 2.594 | 0.1238 |
|  |  |  | PE | F (1, 20) = 0.1095 | 0.7443 |
|  |  |  | PE*AM | F (1, 20) = 0.2307 | 0.6365 |
| **Figure S1b** | EPM (% time open arms) | One-way ANOVA | HU | F (2, 13) = 1.386 | 0.2847 |
| **Figure S1c** | FST (time in immobile behaviour) | One-way ANOVA | HU | F (2, 12) = 0.7908 | 0.4758 |
| **Figure S1d** | EPM (% time open arms) | One-way ANOVA | AM | F (2, 13) = 4.367 | ***0.0328**** |
|  |  |  | AM0.5 |  | 0.1283 |
|  |  |  | AM5 |  | ***0.0428**** |
| **Figure S1e** | FST (time in immobile behaviour) | One-way ANOVA | AM | F (2, 12) = 5.305 | ***0.0224**** |
|  |  |  | AM0.5 |  | ***0.0212**** |
|  |  |  | AM5 |  | 0.0764 |
| **Figure S1f** | NOR (% exploration time) | Student’s  t-tests (paired) | CTRL_f vs CTRL_n | t_7_ = 6.379 | ***0.0004****** |
|  |  |  | HU0.5_f vs HU0.5_n | t_3_ = 3.444 | ***0.0411**** |
|  |  |  | HU5_f vs HU5_n | t_3_ = 4.198 | ***0.0247**** |
| **Figure S1g** | NOR (% exploration time) | Student’s  t-tests (paired) | CTRL_f vs CTRL_n | t_7_ = 6.379 | ***0.0004****** |
|  |  |  | AM0.5_f vs AM0.5_n | t_3_ = 1.412 | 0.2528 |
|  |  |  | AM5_f vs AM5_n | t_3_ = 0.8013 | 0.4815 |
| **Figure S1h1** | Neuronal differentiation (total DCX+ cells) | One-way ANOVA | HU | F (2, 9) = 6.118 | ***0.0210**** |
|  |  |  | HU0.5 |  | 0.4965 |
|  |  |  | HU5 |  | ***0.0140**** |
| **Figure S1h2** | Neuronal differentiation (total BrdU+DCX+ cells) | One-way ANOVA | HU | F (2, 9) = 7.352 | ***0.0128**** |
|  |  |  | HU0.5 |  | ***0.0131**** |
|  |  |  | HU5 |  | ***0.0209**** |
| **Figure S1i1** | Neuronal differentiation (total DCX+ cells) | One-way ANOVA | AM | F (2, 9) = 0.7070 | 0.5186 |
| **Figure S1i2** | FST (time in immobile behaviour) | One-way ANOVA | AM | F (2, 9) = 0.2.203 | 0.1664 |
| **Figure S2c1** | Weight variation | Student’s  t-test | CTRL vs uCMS (unpaired) | t_66_ = 2.731 | ***0.0081***** |
| **Figure S5f** | fEPSP S1/ fEPSP S0 | Student’s  t-test | CTRL vs uCMS (unpaired) | t_4_ = 4.212 | ***0.0136**** |
|  |  | Three-way ANOVA | PE | F (1, 16) = 4.190 | 0.0574 |
|  |  |  | HU vs AM | F (1, 16) = 0.3669 | 0.5532 |
|  |  |  | CB2R treatment | F (1, 16) = 3.911 | 0.0654 |
|  |  |  | PE*HU vs AM | F (1, 16) = 0.1601 | 0.6943 |
|  |  |  | PE*CB2R treatment | F (1, 16) = 3.141 | 0.0954 |
|  |  |  | HU vs AM*CB2R treatment | F (1, 16) = 0.3669 | 0.5532 |
|  |  |  | PE*HU vs AM*CB2R treatment | F (1, 16) = 0.1601 | 0.6943 |
| **Figure S2g** | CB2R expression | Student’s  t-test | CTRL vs uCMS (unpaired) | t_6_ = 2.410 | 0.0526 |
|  |  | Three-way ANOVA | PE | F (1, 24) = 0.1321 | 0.7195 |
|  |  |  | HU vs AM | F (1, 24) = 0.2625 | 0.6131 |
|  |  |  | CB2R treatment | F (1, 24) = 0.7078 | 0.4085 |
|  |  |  | PE*HU vs AM | F (1, 24) = 0.6618 | 0.4239 |
|  |  |  | PE*CB2R treatment | F (1, 24) = 0.09391 | 0.7619 |
|  |  |  | HU vs AM*CB2R treatment | F (1, 24) = 0.2625 | 0.6131 |
|  |  |  | PE*HU vs AM*CB2R treatment | F (1, 24) = 0.6618 | 0.4239 |
| **Figure S3b** | EPM (% time open arms) | Student’s  t-test | CTRL vs uCMS (unpaired) | t_16_ = 3.492 | ***0.0030***** |
|  |  | Two-way ANOVA | HU | F (1, 28) = 2.599 | 0.1182 |
|  |  |  | PE | F (1, 28) = 2.898 | 0.0997 |
|  |  |  | PE*HU | F (1, 28) = 15.64 | ***0.0005****** |
|  |  | Post-hoc | uCMS vs uCMS+PE |  | 0.2851 |
|  |  |  | uCMS vs uCMS+HU+PE |  | 0.0689 |
|  |  |  | uCMS vs uCMS+HU |  | 0.2105 |
| **Figure S3c** | SST (latency) | Student’s  t-test | CTRL vs uCMS (unpaired) | t_16_ = 3.941 | ***0.0012***** |
|  |  | Two-way ANOVA | HU | F (1, 29) = 0.1442 | 0.7069 |
|  |  |  | PE | F (1, 29) = 1.767 | 0.1941 |
|  |  |  | PE*HU | F (1, 29) = 0.01080 | 0.9180 |
| **Figure S3d** | FST (time in immobile behaviour) | Student’s  t-test | CTRL vs uCMS (unpaired) | t_16_ = 6.087 | ***<0.0001 ******* |
|  |  | Two-way ANOVA | HU | F (1, 28) = 1.316 | 0.2611 |
|  |  |  | PE | F (1, 28) = 1.617 | 0.2140 |
|  |  |  | PE*HU | F (1, 28) = 0.6609 | 0.4231 |
| **Figure S3e** | OF (% time in centre) | Student’s  t-test | CTRL vs uCMS (unpaired) | t_15_ = 1.800 | 0.0920 |
|  |  | Two-way ANOVA | HU | F (1, 27) = 0.5928 | 0.4480 |
|  |  |  | PE | F (1, 27) = 0.02890 | 0.8663 |
|  |  |  | PE*HU | F (1, 27) = 0.3913 | 0.5369 |
| **Figure S3f** | NOR (% exploration time) | Student’s  t-tests (paired) | CTRL_f vs CTRL_n | t_8_ = 8.525 | ***<0.0001 ******* |
|  |  |  | uCMS_f vs uCMS_n | t_8_ = 0.8505 | 0.4198 |
|  |  |  | uCMS+PE_f vs uCMS+PE_n | t_6_ = 4.445 | ***0.0044***** |
|  |  |  | uCMS+HU+PE_f vs uCMS+HU+PE_n | t_8_ = 0.7821 | 0.4567 |
|  |  |  | uCMS+HU_f vs uCMS+HU_n | t_6_ = 1.553 | 0.1713 |
| **Figure S4a** | Iba1 occupancy | Student’s  t-test | CTRL vs uCMS (unpaired) | t_4_ = 2.798 | ***0.0489**** |
|  |  | Two-way ANOVA | HU | F (1, 8) = 2.753 | 0.1357 |
|  |  |  | PE | F (1, 8) = 9.275 | 0.0159 |
|  |  |  | PE*HU | F (1, 8) = 1.894 | 0.2060 |
|  |  | Post-hoc | uCMS vs uCMS+PE |  | ***0.0348**** |
|  |  |  | uCMS vs uCMS+HU+PE |  | ***0.0260**** |
|  |  |  | uCMS vs uCMS+HU |  | 0.1480 |
| **Figure S4b** | MBP occupancy | Student’s  t-test | CTRL vs uCMS (unpaired) | t_4_ = 30.03 | ***<0.0001 ******* |
|  |  | Two-way ANOVA | HU | F (1, 8) = 4.903 | 0.0577 |
|  |  |  | PE | F (1, 8) = 0.04151 | 0.8437 |
|  |  |  | PE*HU | F (1, 8) = 1.663 | 0.2332 |
| **Figure S4c** | GFAP occupancy | Student’s  t-test | CTRL vs uCMS (unpaired) | t_4_ = 1.302 | 0.2627 |
|  |  | Two-way ANOVA | HU | F (1, 8) = 0.2501 | 0.6304 |
|  |  |  | PE | F (1, 8) = 0.0006983 | 0.9796 |
|  |  |  | PE*HU | F (1, 8) = 0.003476 | 0.9544 |
| **Figure S4d** | TNFα expression | Student’s  t-test | CTRL vs uCMS (unpaired) | t_11_ = 1.456 | 0.1732 |
|  |  | Two-way ANOVA | HU | F (1, 21) = 0.1640 | 0.6896 |
|  |  |  | PE | F (1, 21) = 2.147 | 0.1576 |
|  |  |  | PE*HU | F (1, 21) = 6.413 | ***0.0194**** |
|  |  | Post-hoc | uCMS vs uCMS+PE |  | ***0.0296**** |
|  |  |  | uCMS vs uCMS+HU+PE |  | 0.4371 |
|  |  |  | uCMS vs uCMS+HU |  | 0.1132 |
| **Figure S4e** | IL-1β expression | Student’s  t-test | CTRL vs uCMS (unpaired) | t_10_ = 2.165 | 0.0557 |
|  |  | Two-way ANOVA | HU | F (1, 20) = 2.242 | 0.1499 |
|  |  |  | PE | F (1, 20) = 0.09377 | 0.7626 |
|  |  |  | PE*HU | F (1, 20) = 0.6887 | 0.4164 |
| **Figure S4f** | IL-10 expression | Student’s  t-test | CTRL vs uCMS (unpaired) | t_10_ = 0.7400 | 0.4763 |
|  |  | Two-way ANOVA | HU | F (1, 20) = 1.039 | 0.3203 |
|  |  |  | PE | F (1, 20) = 0.4504 | 0.5098 |
|  |  |  | PE*HU | F (1, 20) = 1.782 | 0.1969 |
| **Figure S4g** | Arginase expression | Student’s  t-test | CTRL vs uCMS (unpaired) | t_11_ = 1.302 | 0.2196 |
|  |  | Two-way ANOVA | HU | F (1, 20) = 0.8134 | 0.3778 |
|  |  |  | PE | F (1, 20) = 0.1501 | 0.7025 |
|  |  |  | PE*HU | F (1, 20) = 0.2680 | 0.6103 |
| **Figure S4h** | iNOS expression | Student’s  t-test | CTRL vs uCMS (unpaired) | t_11_ = 1.070 | 0.3075 |
|  |  | Two-way ANOVA | HU | F (1, 20) = 0.2389 | 0.6304 |
|  |  |  | PE | F (1, 20) = 0.1469 | 0.7056 |
|  |  |  | PE*HU | F (1, 20) = 0.06651 | 0.7991 |
| **Figure S5b** | fEPSP S1/ fEPSP S0 | Student’s  t-test | CTRL vs uCMS (unpaired) | t_4_ = 4.212 | ***0.0136**** |
|  |  | Three-way ANOVA | PE | F (1, 16) = 4.190 | 0.0574 |
|  |  |  | HU vs AM | F (1, 16) = 0.3669 | 0.5532 |
|  |  |  | CB2R treatment | F (1, 16) = 3.911 | 0.0654 |
|  |  |  | PE*HU vs AM | F (1, 16) = 0.1601 | 0.6943 |
|  |  |  | PE*CB2R treatment | F (1, 16) = 3.141 | 0.0954 |
|  |  |  | HU vs AM*CB2R treatment | F (1, 16) = 0.3669 | 0.5532 |
|  |  |  | PE*HU vs AM*CB2R treatment | F (1, 16) = 0.1601 | 0.6943 |
| **Figure S5e** | CB2R expression | Student’s  t-test | CTRL vs uCMS (unpaired) | t_6_ = 2.410 | 0.0526 |
|  |  | Three-way ANOVA | PE | F (1, 24) = 0.1321 | 0.7195 |
|  |  |  | HU vs AM | F (1, 24) = 0.2625 | 0.6131 |
|  |  |  | CB2R treatment | F (1, 24) = 0.7078 | 0.4085 |
|  |  |  | PE*HU vs AM | F (1, 24) = 0.6618 | 0.4239 |
|  |  |  | PE*CB2R treatment | F (1, 24) = 0.09391 | 0.7619 |
|  |  |  | HU vs AM*CB2R treatment | F (1, 24) = 0.2625 | 0.6131 |
|  |  |  | PE*HU vs AM*CB2R treatment | F (1, 24) = 0.6618 | 0.4239 |
| **Table 1**  **(Ki67+ cells)** | Dorsal: Supra SGZ | Student’s  t-test | CTRL vs uCMS (unpaired) | t_8_ = 1.504 | 0.1710 |
|  |  | Two-way ANOVA | AM | F (1, 13) = 0.02995 | 0.8653 |
|  |  |  | PE | F (1, 13) = 0.02219 | 0.8839 |
|  |  |  | PE*AM | F (1, 13) = 0.3730 | 0.5519 |
|  | Dorsal: Infra SGZ | Student’s  t-test | CTRL vs uCMS (unpaired) | t_8_ = 0.1372 | 0.8943 |
|  |  | Two-way ANOVA | AM | F (1, 13) = 0.03512 | 0.8542 |
|  |  |  | PE | F (1, 13) = 0.04826 | 0.8295 |
|  |  |  | PE*AM | F (1, 13) = 0.002391 | 0.9617 |
|  | Dorsal: Supra GCL | Student’s  t-test | CTRL vs uCMS (unpaired) | t_8_ = 9.113 | ***<0.0001 ******* |
|  |  | Two-way ANOVA | AM | F (1, 13) = 5.113 | ***0.0415**** |
|  |  |  | PE | F (1, 13) = 10.40 | ***0.0066***** |
|  |  |  | PE*AM | F (1, 13) = 0.08281 | 0.7781 |
|  |  | Post-hoc | uCMS vs uCMS+PE |  | 0.0706 |
|  |  |  | uCMS vs uCMS+AM+PE |  | ***0.0047***** |
|  |  |  | uCMS vs uCMS+AM |  | 0.2387 |
|  | Dorsal: Infra GCL | Student’s  t-test | CTRL vs uCMS (unpaired) | t_8_ = 3.458 | ***0.0086***** |
|  |  | Two-way ANOVA | AM | F (1, 13) = 0.8239 | 0.3806 |
|  |  |  | PE | F (1, 13) = 0.5147 | 0.4858 |
|  |  |  | PE*AM | F (1, 13) = 2.263 | 0.1564 |
|  | Ventral: Supra SGZ | Student’s  t-test | CTRL vs uCMS (unpaired) | t_8_ = 3.245 | ***0.0118**** |
|  |  | Two-way ANOVA | AM | F (1, 13) = 0.06641 | 0.8007 |
|  |  |  | PE | F (1, 13) = 1.345 | 0.2671 |
|  |  |  | PE*AM | F (1, 13) = 0.5312 | 0.4790 |
|  | Ventral: Infra SGZ | Student’s  t-test | CTRL vs uCMS (unpaired) | t_8_ = 0.09911 | 0.9235 |
|  |  | Two-way ANOVA | AM | F (1, 13) = 0.8533 | 0.3724 |
|  |  |  | PE | F (1, 13) = 0.1207 | 0.7338 |
|  |  |  | PE*AM | F (1, 13) = 1.435 | 0.2523 |
|  | Ventral: Supra GCL | Student’s  t-test | CTRL vs uCMS (unpaired) | t_8_ = 2.874 | ***0.0207**** |
|  |  | Two-way ANOVA | AM | F (1, 13) = 0.009690 | 0.9231 |
|  |  |  | PE | F (1, 13) = 7.010 | ***0.0201**** |
|  |  |  | PE*AM | F (1, 13) = 0.4116 | 0.5323 |
|  |  | Post-hoc | uCMS vs uCMS+PE |  | 0.0950 |
|  |  |  | uCMS vs uCMS+AM+PE |  | 0.2386 |
|  |  |  | uCMS vs uCMS+AM |  | 0.9729 |
|  | Ventral: Infra GCL | Student’s  t-test | CTRL vs uCMS (unpaired) | t_8_ = 1.487 | 0.1752 |
|  |  | Two-way ANOVA | AM | F (1, 13) = 0.7912 | 0.3899 |
|  |  |  | PE | F (1, 13) = 0.3127 | 0.5855 |
|  |  |  | PE*AM | F (1, 13) = 0.7533 | 0.4012 |
| **Table 1**  **(BrdU+NeuN+ cells)** | Dorsal: Supra SGZ | Student’s  t-test | CTRL vs uCMS (unpaired) | t_8_ = 7.732 | ***<0.0001 ******* |
|  |  | Two-way ANOVA | AM | F (1, 13) = 2.986 | 0.1077 |
|  |  |  | PE | F (1, 13) = 52.55 | ***<0.0001 ******* |
|  |  |  | PE*AM | F (1, 13) = 0.03255 | 0.8596 |
|  |  | Post-hoc | uCMS vs uCMS+PE |  | ***0.0006****** |
|  |  |  | uCMS vs uCMS+AM+PE |  | ***<0.0001 ******* |
|  |  |  | uCMS vs uCMS+AM |  | 0.6287 |
|  | Dorsal: Infra SGZ | Student’s  t-test | CTRL vs uCMS (unpaired) | t_8_ = 0.4664 | 0.6534 |
|  |  | Two-way ANOVA | AM | F (1, 13) = 0.2517 | 0.6206 |
|  |  |  | PE | F (1, 13) = 0.5663 | 0.4651 |
|  |  |  | PE*AM | F (1, 13) = 0.1426 | 0.7118 |
|  | Dorsal: Supra GCL | Student’s  t-test | CTRL vs uCMS (unpaired) | t_8_ = 5.039 | ***0.0010***** |
|  |  | Two-way ANOVA | AM | F (1, 13) = 0.01751 | 0.8967 |
|  |  |  | PE | F (1, 13) = 29.59 | ***0.0001****** |
|  |  |  | PE*AM | F (1, 13) = 1.182 | 0.2966 |
|  |  | Post-hoc | uCMS vs uCMS+PE |  | ***0.0223**** |
|  |  |  | uCMS vs uCMS+AM+PE |  | ***0.0059***** |
|  |  |  | uCMS vs uCMS+AM |  | 0.7748 |
|  | Dorsal: Infra GCL | Student’s  t-test | CTRL vs uCMS (unpaired) | t_8_ = 4.084 | ***0.0035***** |
|  |  | Two-way ANOVA | AM | F (1, 13) = 12.88 | ***0.0033***** |
|  |  |  | PE | F (1, 13) = 11.17 | ***0.0057***** |
|  |  |  | PE*AM | F (1, 13) = 2.570 | 0.1329 |
|  |  | Post-hoc | uCMS vs uCMS+PE |  | ***0.0098***** |
|  |  |  | uCMS vs uCMS+AM+PE |  | ***0.0007****** |
|  |  |  | uCMS vs uCMS+AM |  | ***0.0070***** |
|  | Ventral: Supra SGZ | Student’s  t-test | CTRL vs uCMS (unpaired) | t_8_ = 3.934 | ***0.0043***** |
|  |  | Two-way ANOVA | AM | F (1, 13) = 10.65 | ***0.0062***** |
|  |  |  | PE | F (1, 13) = 6.394 | ***0.0252**** |
|  |  |  | PE*AM | F (1, 13) = 0.1797 | 0.6786 |
|  |  | Post-hoc | uCMS vs uCMS+PE |  | 0.3863 |
|  |  |  | uCMS vs uCMS+AM+PE |  | ***0.0031***** |
|  |  |  | uCMS vs uCMS+AM |  | 0.1686 |
|  | Ventral: Infra SGZ | Student’s  t-test | CTRL vs uCMS (unpaired) | t_8_ = 2.452 | ***0.0468**** |
|  |  | Two-way ANOVA | AM | F (1, 13) = 2.899 | 0.1124 |
|  |  |  | PE | F (1, 13) = 0.8157 | 0.3829 |
|  |  |  | PE*AM | F (1, 13) = 4.826 | ***0.0468**** |
|  |  | Post-hoc | uCMS vs uCMS+PE |  | 0.1214 |
|  |  |  | uCMS vs uCMS+AM+PE |  | 0.2234 |
|  |  |  | uCMS vs uCMS+AM |  | ***0.0417**** |
|  | Ventral: Supra GCL | Student’s  t-test | CTRL vs uCMS (unpaired) | t_8_ = 0.1810 | 0.8609 |
|  |  | Two-way ANOVA | AM | F (1, 13) = 2.232 | 0.1590 |
|  |  |  | PE | F (1, 13) = 0.003285 | 0.9552 |
|  |  |  | PE*AM | F (1, 13) = 0.4174 | 0.5295 |
|  | Ventral: Infra GCL | Student’s  t-test | CTRL vs uCMS (unpaired) | t_8_ = 0.9618 | 0.3643 |
|  |  | Two-way ANOVA | AM | F (1, 13) = 0.5910 | 0.4558 |
|  |  |  | PE | F (1, 13) = 6.511 | ***0.0241**** |
|  |  |  | PE*AM | F (1, 13) = 5.441 | ***0.0364**** |
|  |  | Post-hoc | uCMS vs uCMS+PE |  | 0.9981 |
|  |  |  | uCMS vs uCMS+AM+PE |  | 0.0912 |
|  |  |  | uCMS vs uCMS+AM |  | 0.6211 |
